# LRRK2 mediates haloperidol-induced changes in indirect pathway striatal projection neurons

**DOI:** 10.1101/2024.06.06.597594

**Authors:** Chuyu Chen, Meghan Masotti, Nathaniel Shepard, Vanessa Promes, Giulia Tombesi, Daniel Arango, Claudia Manzoni, Elisa Greggio, Sabine Hilfiker, Yevgenia Kozorovitskiy, Loukia Parisiadou

## Abstract

Haloperidol is used to manage psychotic symptoms in several neurological disorders through mechanisms that involve antagonism of dopamine D2 receptors that are highly expressed in the striatum. Significant side effects of haloperidol, known as extrapyramidal symptoms, lead to motor deficits similar to those seen in Parkinson’s disease and present a major challenge in clinical settings. The underlying molecular mechanisms responsible for these side effects remain poorly understood. Parkinson’s disease-associated LRRK2 kinase has an important role in striatal physiology and a known link to dopamine D2 receptor signaling. Here, we systematically explore convergent signaling of haloperidol and LRRK2 through pharmacological or genetic inhibition of LRRK2 kinase, as well as knock-in mouse models expressing pathogenic mutant LRRK2 with increased kinase activity. Behavioral assays show that LRRK2 kinase inhibition ameliorates haloperidol-induced motor changes in mice. A combination of electrophysiological and anatomical approaches reveals that LRRK2 kinase inhibition interferes with haloperidol-induced changes, specifically in striatal neurons of the indirect pathway. Proteomic studies and targeted intracellular pathway analyses demonstrate that haloperidol induces a similar pattern of intracellular signaling as increased LRRK2 kinase activity. Our study suggests that LRRK2 kinase plays a key role in striatal dopamine D2 receptor signaling underlying the undesirable motor side effects of haloperidol. This work opens up new therapeutic avenues for dopamine-related disorders, such as psychosis, also furthering our understanding of Parkinson’s disease pathophysiology.

**Summary:** Chen et al. demonstrate that haloperidol mediated changes in the striatal indirect pathway neurons and circuits are linked to Parkinson’s disease associated LRRK2. Inhibiting LRRK2 kinase activity ameliorates the motoric side effects of haloperidol, suggesting a potential approach to alleviating the unwanted side effects of antipsychotics.

## Introduction

Dopamine D2 receptors (D2Rs) are important pharmacological targets for neuropsychiatric diseases such as schizophrenia and Parkinson’s disease (PD)^1,2^. D2Rs are highly expressed in spiny projection neurons (SPNs) of the indirect striatal pathway (iSPNs), one of striatum’s two principal and functionally neuronal subtypes—indirect striatal pathway, together with the direct striatal pathway, controls movement^3,4,5,6^. D2Rs inhibit cAMP production by coupling with Gi/o and, as a result, reduce protein kinase A (PKA) sinaling^7^. These intracellular events mediated by D2R signaling pathways have a significant impact on the physiological functions of iSPNs at both cell-intrinsic and network levels^8,9,10^.

D2Rs are known targets of all antipsychotics^11^. Haloperidol, an antagonist of D2Rs, is a first-generation typical antipsychotic that is still widely used to treat various neuropsychiatric diseases^12^. However, it can cause major side effects, including severe motor deficits that mimic PD symptoms, collectively known as extrapyramidal motor symptoms^13,14^. In rodents, haloperidol administration induces catalepsy, a state of inhibited movement that results from the blockade of D2 receptors in the nigrostriatal pathway^15,16^. While the side effect profile of this otherwise effective antipsychotic presents a significant challenge in the clinic, details of the signaling cascades and neural circuit adaptations that lead to haloperidol-induced motor deficits remain elusive. Developing efficient and safer therapies for treating D2R-related disorders remains challenging due to the heterogeneous expression of D2R across cell types and diverse cascades downstream of D2R.

PD is a disorder characterized by the loss of the nigrostriatal dopamine neurons, which results in significant motor deficits. Variants of several genes have been linked to familial PD, with the kinase LRRK2 being particularly interesting due to its association with both familial and idiopathic PD^17,18,19^. Previous studies have found that, in the striatum, LRRK2 is highly expressed in SPNs and plays a crucial role in regulating the striatal function ^20,22,22,23,24,25,26^ including via modulation of D2R signaling^27–29^. Pathogenic PD-associated LRRK2 mutations lead to kinase hyperactivity^30,31^, suggesting that LRRK2 activity is important in the pathophysiological mechanisms of PD. Thus, animal models carrying human-disease-linked LRRK2 mutations can be used as a genetic proxy to understand the consequences of enhanced kinase activity on striatal neuronal signaling, structure, and function.

The most etiologically relevant knock-in (KI) mice expressing pathogenic LRRK2 mutations exhibit either normal or moderately altered motor behaviors^32–34^. However, perturbation of D2R signaling in these animals unmasks motor phenotypes. Impairment of striatal motor learning as a result of D2R antagonism is enhanced by LRRK2 mutations^23^. Conversely, impairment of locomotor activity by activation of D2R signaling is partially ameliorated in LRRK2 KI mice ^34^. Although LRRK2 activity and D2R signaling appear as convergent themes across different studies and genetic models, a coherent synthesis of these results, accounting for variability in model design, D2R manipulation, and technical approaches, is currently lacking.

The current study aims to understand how LRRK2 modulates the effects of haloperidol on the behavioral, physiological, and molecular changes downstream of D2R antagonism in the striatum. To achieve this, we use a combination of behavioral tests, whole-cell electrophysiology, proteomic and phosphoproteomic analyses as well as transcript level measurements in concert with pharmacological and genetic manipulations of LRRK2 activity. Our research demonstrates a critical role for LRRK2 kinase activity in haloperidol-induced motor changes. We find that inhibition of LRRK2 kinase activity can alleviate these motor side effects, paralleled by a reversal of both structural and functional indirect pathway changes induced by haloperidol. Conversely, LRRK2 hyperactivity, the result of PD-linked pathogenic mutations, mimics the molecular and biochemical effects of haloperidol on striatal intracellular signaling. Overall, our work opens new directions for the therapeutic management of psychosis and helps to refine the pathological mechanisms relevant to PD.

## Results

### LRRK2 kinase activity mediates the effects of haloperidol on movement disruption

In order to examine the link between LRRK2 activity and motor behaviors sensitive to D2R signaling, we evaluated whether the effects of blocking D2R signaling by haloperidol are influenced by LRRK2 inhibition. Haloperidol, a typical antipsychotic used to treat psychosis, is known to cause Parkinsonian-like side effects, including catalepsy in rodents and dyskinesia in humans^14^. We hypothesized that if LRRK2 kinase activity plays a significant role in D2R signaling in the striatum, its inhibition may interfere with the cataleptic response induced by haloperidol.

Wild-type (WT) mice were injected with a subchronic dose (1 mg/kg) of haloperidol or vehicle control for 7 days, followed by the measurement of their cataleptic response^15^. A potent and selective LRRK2 kinase inhibitor MLi-2^35, 36, 37^ was administered at 10 mg/kg daily^38^ 30 mins before haloperidol (Fig. 1A). Consistent with previous studies^16^, seven days of haloperidol administration without LRRK2 inhibition induced a significant cataleptic response in WT mice, as measured by the bar test (p<0.0001, Tukey’s multiple comparisons test after one-way ANOVA; n=9-15 mice). MLi-2 itself had no effect on catalepsy compared to vehicle control (p>0.999 post-hocs, as noted, n=8-9 mice). However, when haloperidol and MLi-2 were combined, the haloperidol-induced cataleptic response was significantly attenuated (p=0.0030, Tukey’s multiple comparisons post-hoc after one-way ANOVA as noted, n=13-15 mice) (Fig. 1B). In order to confirm that this MLi-2 administration was sufficient to decrease LRRK2 kinase activity in vivo, we measured phosphorylation of the Ser106 site of Rab12, a well-established measure of LRRK2 kinase activity^38,39^. Consistent with reduced LRRK2 activity, MLi-2 administration resulted in decreased phosphorylation of Rab12 at Ser106, compared to vehicle controls (p=0.0043 Tukey’s multiple comparisons test after one-way ANOVA; n=6 mice (Suppl. Fig. 1A-B). Total Rab12 levels were found unaltered across treatments (p=0.9994, Tukey’s multiple comparisons test after one-way ANOVA, Suppl. Fig. 1C).

**Figure 1:**
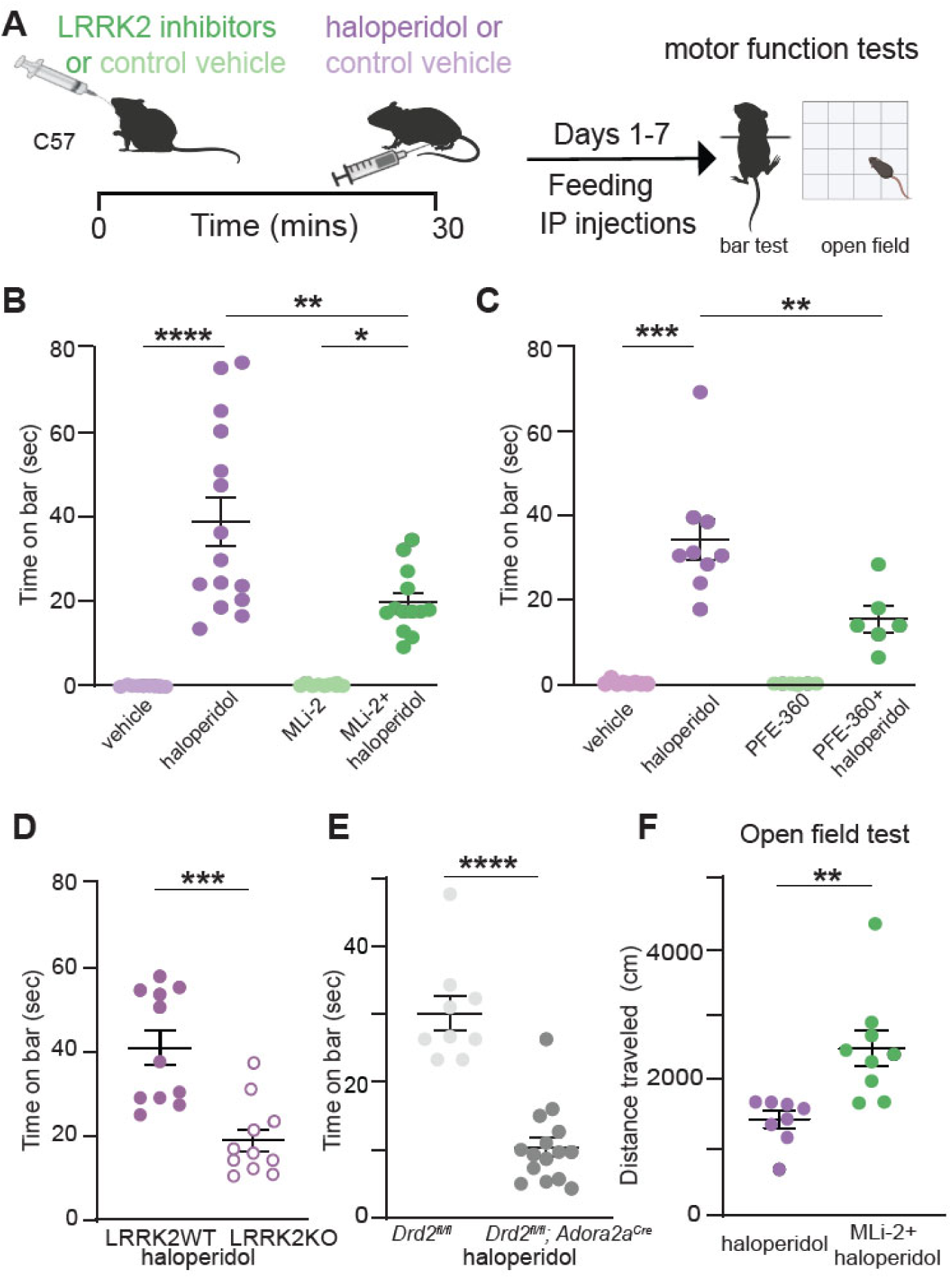
LRRK2 mediates the effects of haloperidol on movement disruption. **A.** Schematic of the experiment and treatment schedule. Catalepsy was assessed with the bar test, and distance traveled was recorded in an open-field arena; both were performed 1 hr after the final treatment administration. Treatment conditions: Vehicle for MLi-2, 40% 2-hydroxypropyl-β-cyclodextrin; MLi2, 10 mg/kg; vehicle for PFE-360, 1.25% hydroxypropyl cellulose + 0.5% docusate sodium; PFE-360, 5 mg/kg; vehicle for haloperidol, saline (0.9% NaCl); haloperidol, 1 mg/kg. Some schematics were created with BioRender.com. X axis: vehicle = vehicle for MLi-2+ vehicle for haloperidol; haloperidol = vehicle for MLi-2+haloperidol; MLi-2 or PFE-360 = MLi-2 or PFE-360+vehicle for haloperidol; MLi-2 or PFE-360+haloperidol, as indicated. All drugs were administered intraperitoneally (i.p.) except MLi-2, PFE-360, and their vehicles, which were given orally. **B.** Cataleptic response of WT mice receiving haloperidol, MLi-2, their combination, and vehicle controls for 7 days. N=9, 15, 8, 13 mice, in order of groups presented. **C.** Same as B, but using PFE-360, and its corresponding vehicle. N=8, 9, 6, 6 mice, in order of groups presented. **D.** Cataleptic response of LRRK2-WT and LRRK2-KO mice receiving haloperidol or control vehicle for 7 days. N=11 mice for both genotypes. **E.** Cataleptic response of *Drd2*^fl/fl^ and *Drd2^fl/fl^; Adora2a^cre^* mice receiving haloperidol for 7 days. N=9 and 15 mice, respectively. **F.** Distance traveled in the open-field test for WT mice receiving haloperidol or haloperidol + MLi-2. N=8 and 9 mice, respectively. Data are represented as mean±SEM. Asterisks in B-C show statistical significance for Tukey’s multiple comparison tests after one-way ANOVA; *p < 0.05, **p< 0.01, ***p < 0.001, ****p < 0.0001. Asterisks in D-F show statistical significance for unpaired t-test comparison; **p< 0.01,***p < 0.001, ****p < 0.0001.

To ensure that the observed effects were not limited to the pharmacology of the MLi-2 inhibitor, we treated mice with another potent and specific LRRK2 kinase inhibitor, PFE-360^40,41^ following the same treatment schedule (Fig. 1A). PFE-360 and MLi-2 have distinct structures and off-target profiles. PFE-360, like MLI-2, also reduced haloperidol-induced catalepsy in WT mice (p=0.0024, Tukey’s multiple comparisons post-hoc after one-way ANOVA, n=6-9 mice) (Fig. 1C). To further verify the relevance of LRRK2 kinase activity for haloperidol-induced catalepsy, we examined the effect of genetic ablation of LRRK2 on the motor side effects of haloperidol. Consistent with our pharmacological inhibition data, we found that catalepsy in haloperidol-treated LRRK2-KO mice was significantly lower than that in LRRK2-WT mice (p=0.0002, unpaired t-test, n= 11 mice) (Fig. 1D).

Haloperidol has a strong affinity for D2Rs, but it lacks selectivity^42^. To determine if haloperidol-associated catalepsy is primarily the result of inhibition of D2R signaling in striatal iSPNs, we generated a mouse line where *Drd2*, which encodes the D2R, was selectively deleted in iSPNs (*Drd2^fl/fl^*). The cataleptic response of *Drd2^fl/fl^* mice was significantly reduced compared to littermate controls (p<0.0001, unpaired t-test, n=9-15) (Fig. 1E). These data suggest that LRRK2 contributes to the induction of catalepsy as a result of blockade of D2R signaling in iSPNs.

Next, to investigate if LRRK2 inhibition modulates haloperidol effects in a different motor behavior task, we assessed locomotor behavior of mice in an open field arena after treatment with haloperidol and MLi-2. WT mice were treated for seven days with haloperidol, or haloperidol and MLi-2, with an open field test one hour after the final treatment. Treatment with haloperidol significantly reduced the total distance traveled (p<0.0001, Tukey’s multiple comparisons test after one way ANOVA, n=8 mice), while MLi-2 alone had no effect on locomotion (p=0.8803, Tukey’s multiple comparisons test after one-way ANOVA, n=7-8) (Suppl. Fig. 1D). Co-administration of MLi-2 and haloperidol increased the distance traveled, compared to haloperidol alone (p=0.0039, unpaired t-test, n=8-9 mice, Fig. 1F). These findings suggest that motor behavior changes induced by subchronic haloperidol administration broadly involve LRRK2.

To better understand the time course of these behavioral effects, we tested the impact of LRRK2 kinase inhibitors on catalepsy after acute (60 mins) and chronic (14 days) 1 mg/kg haloperidol administration. In both cases, WT mice were given combined treatments with MLi-2 or vehicle controls 30 min before haloperidol. Catalepsy was measured 1 hr after the last injection (Suppl. Fig. 1E). In line with previous studies, acute administration of haloperidol (1 mg/kg, 1 hr) induced catalepsy^43,15,44^ as measured with a standard bar test, compared to vehicle controls (p<0.0001, Tukey’s multiple comparisons post-hoc after one-way ANOVA, n=5-15 mice) (Suppl. Fig. 1F). Co-administration of haloperidol and MLi-2 decreased the cataleptic response compared to haloperidol-treated mice (p=0.021, Tukey’s multiple comparisons post-hoc after one-way ANOVA, n=12-15 mice) (Suppl. Fig 1F). This timing and dosing of MLi-2 co-administration was also sufficient to decrease the phosphorylation of LRRK2 downstream target p-Rab12^38^ (Suppl Fig. 1H). In the chronic paradigm, we found that MLi-2 had no effect on haloperidol-induced catalepsy after 14 days (p=0.7328, Tukey’s multiple comparisons post-hoc after one-way ANOVA, n=14-16 mice) (Suppl. Fig 1G). This is likely due to developing tolerance to the cataleptic effects of the drug^45,16^. Thus, MLi-2 is no longer effective at modulating the substantially declined cataleptic response (Suppl. Fig. 1I). Taken together, these results are consistent with a critical role of LRRK2 kinase in the events that underlie the motor disruption induced by D2R antagonism. Finally, no sex differences were found in treatment paradigms (Suppl. Fig. 2).

Several studies highlight the importance of dopamine D2R signaling in striatal motor learning^46,47, 48^, and our previous data report differences with LRRK2 mutations in the presence of D2R antagonists in striatal motor learning^23^. These observations suggest that LRRK2 may interfere with D2R signaling during skill acquisition. Therefore, we determined the impact of LRRK2 kinase activity on modulating the effects of haloperidol using the accelerating rotarod test. We assessed the latency to fall in WT mice treated with either haloperidol, MLi-2, or their controls for 14 days in an accelerated rotarod (Suppl. Fig.3A). The average latency to fall improved towards the end of the task in the control-treated mice, treatment factor F3,344=22.35, p<0.0001, Session Factor F7,344=7.553, p<0.0001. Treatment x session interaction F21,344=0.8274, p=0.6857, Tukey’s multiple comparisons test after two-way ANOVA, vehicle, n=10; haloperidol, n=10; MLi2, n=16; MLi2+ haloperidol, n=11. (Suppl. Fig. 3B) Haloperidol significantly decreased the latency to fall, compared to vehicle controls over the last three sessions of the test (p<0.001, Tukey’s multiple comparisons post-hoc after one-way ANOVA). This decrease was prevented in mice co-administered the LRRK2 kinase inhibitor MLi-2 (p=0.0011, Tukey’s multiple comparisons after one way ANOVA) (Suppl. Fig. 3C). Our observations suggest that LRRK2 interferes with haloperidol-mediated effects on striatal motor learning.

### LRRK2 kinase activity regulates haloperidol-induced adaptations in striatal indirect pathway circuits

Haloperidol is reported to induce a series of homeostatic synaptic and intrinsic adaptations primarily in the indirect pathway circuits^16,49–52^. These adaptations include dendritic spine loss in iSPNs, which occurs after 5 days and persists for up to 14 days of daily treatment (Fig. 2A) (synaptic adaptations). Additionally, a decrease in the excitability of iSPNs has been observed after 14 days of haloperidol administration, which is thought to reflect an intrinsic homeostatic change aimed at normalizing the persistent increase in firing rate after D2R antagonism (Fig. 2A) (intrinsic adaptations)^16,53^. The cataleptic tolerance (e.g., decreased response to haloperidol over time)^45^ between sub-chronic (5 days) and chronic (14 days) haloperidol treatment, described above, has been suggested to reflect remodeling of the indirect pathway circuits^16^. However, the exact intrinsic and network changes underlying these processes remain largely unknown.

**Figure 2.**
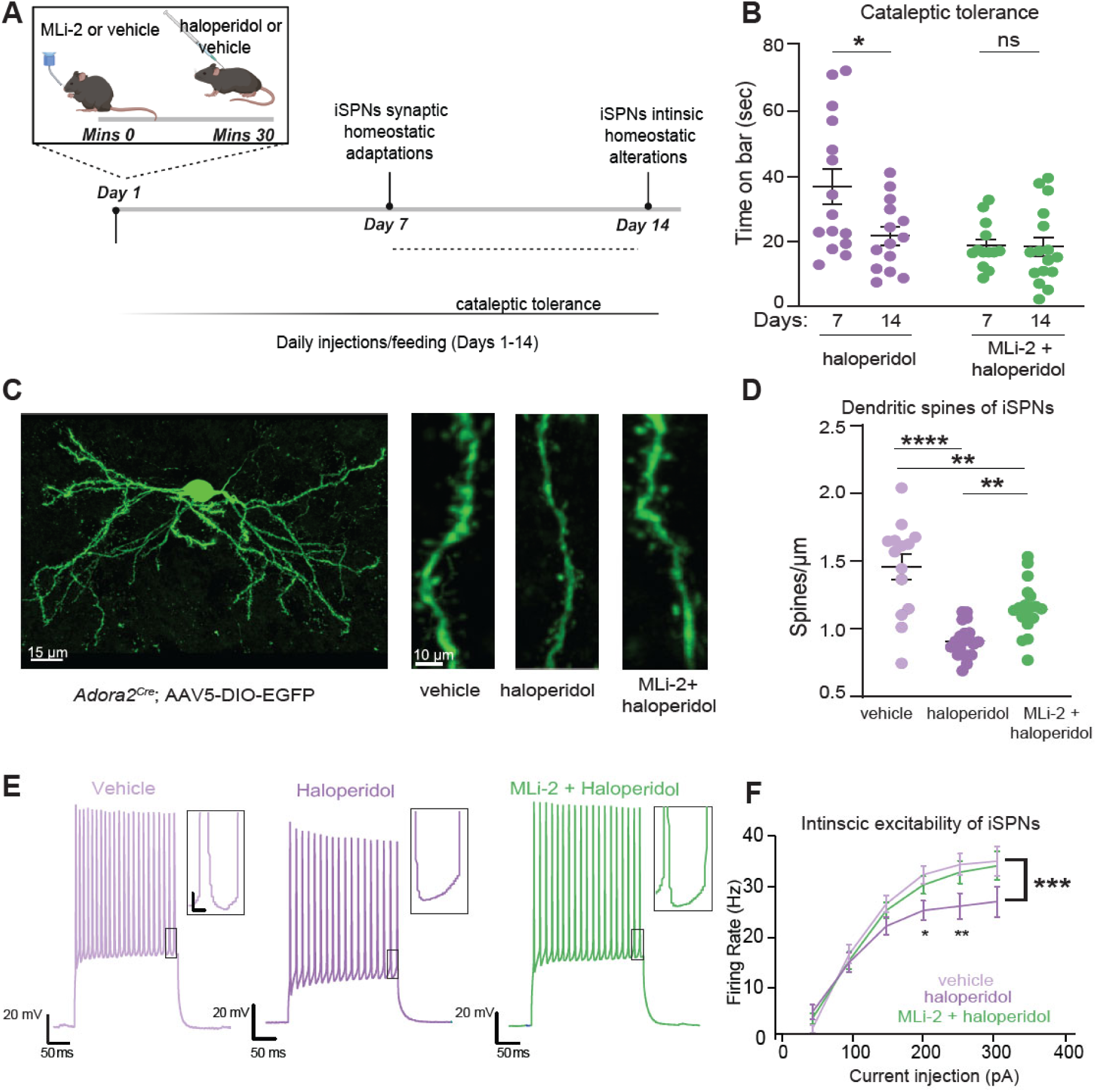
LRRK2 interferes with haloperidol induced adaptations in indirect pathway SPNs. **A.** Schematic of the proposed timeline for synaptic and cell-intrinsic adaptations with haloperidol treatment, based on a synthesis of current and prior findings. Some of the schematics were created with BioRender.com. **B.** Cataleptic response of WT mice receiving haloperidol, or haloperidol + MLi-2, at 7 and 14 days following treatments. Haloperidol-treated mice develop tolerance to catalepsy after 14 but not 7 days of haloperidol administration. N=15, 14, 13, 16 mice, in order of groups presented. Data for 7- and 14-day regimens are from panels 1B and Suppl. Fig. 1G, respectively. **C.** *Left,* Example of confocal maximum projection image of an *Adora2a^Cre^* SPN expressing AAV5/DIO-EGFP. Scale bar=15 μm. *Right,* Examples of dendritic segments in different treatment conditions. Scale bar=10 μm. **D.** Summary graph of dendritic spine density in iSPNs across treatments. N=14-20 cells, 3 mice/group. **E.** Example current clamp recording traces in response to a 150 pA current injection in iSPNs of WT mice after 14 days of treatment, as noted. Recordings were performed 24-48 hours after the final injection. Insets: scale 4 mV, 20 ms. **F.** Summary data show decreased cellular excitability with haloperidol, rescued to control levels by LRRK2 inhibition. Scale, as noted. N=14-29 cells, 4-6 mice/condition. Data are represented as mean±SEM. Asterisks in B show statistical significance for Tukey’s multiple comparison tests after two-way ANOVA; *p< 0.05. D shows statistical significance for Tukey’s multiple comparison tests after one-way ANOVA**p< 0.01, ****, p<0.0001. Asterisks in F show statistical significance after two-way ANOVA with multiple comparisons. Large asterisk, interaction between treatment and current steps. Small asterisks, significance at specific current steps, determined by Sidak multiple comparisons. *p<0.05.

**Figure 3:**
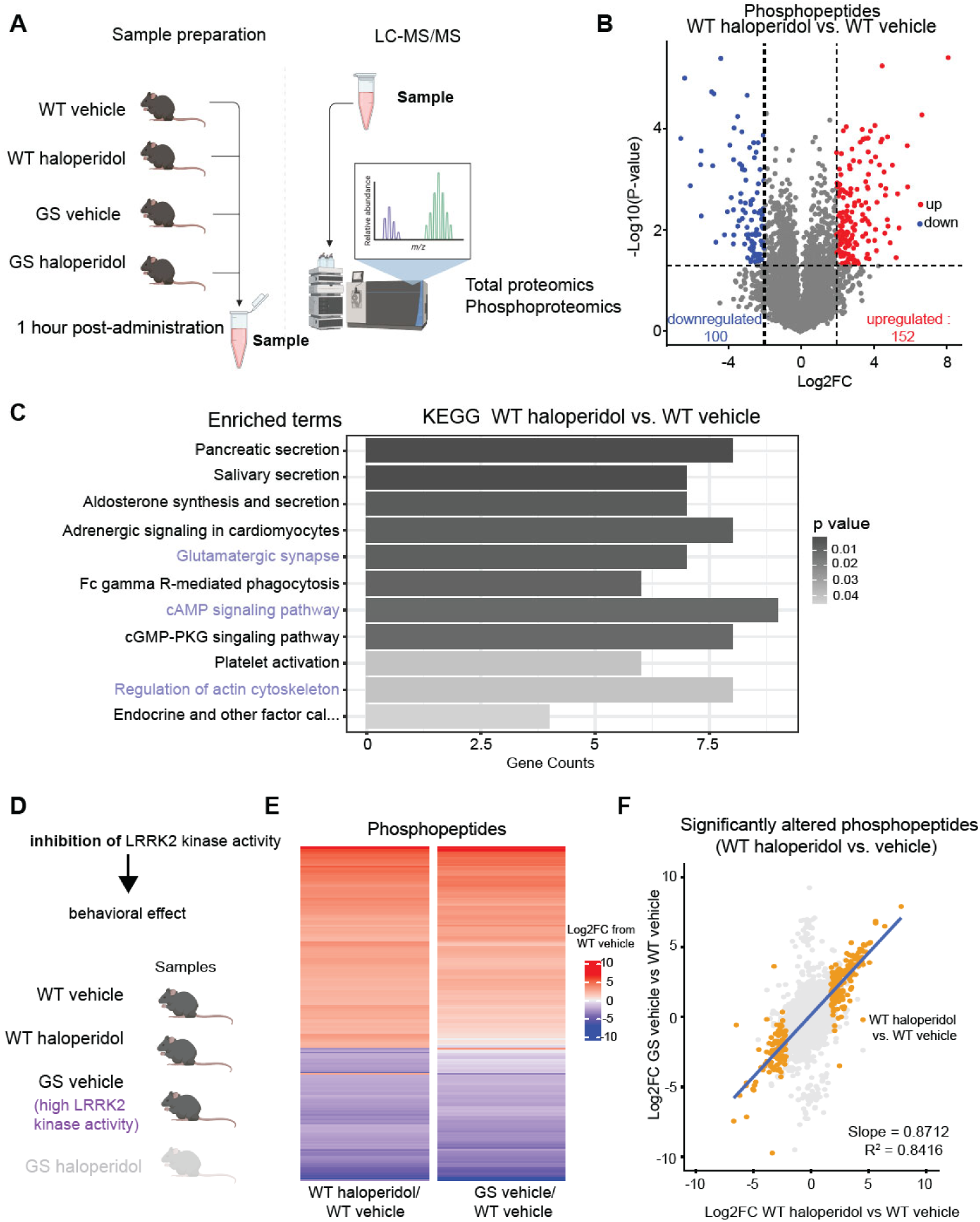
Haloperidol treatment induces similar changes in the striatal phosphoproteome as increased LRRK2 kinase activity. **A.** Schematic of experimental procedures. WT and GS mutant mice were treated with haloperidol or vehicle, and striatal samples were harvested after 1 hour. LC-MS/MS analyses for total proteins and phosphopeptides were conducted on the same striatal samples. N=3 mice/group. Parts of the schematic were created with BioRender.com. **B.** Volcano plot comparing the phosphopeptides between haloperidol and vehicle-treated WT mice. Phosphopeptides that are significantly differentially regulated (|Log2FC| > 2 and unadjusted p-value ≤ 0.05 by multiple unpaired t-tests) are colored red and blue for up- and down-regulated, respectively. **C.** Results of Gene Set Enrichment Analysis (GSEA) of the KEGG gene set for genes with at least one differentially regulated phosphopeptide in WT haloperidol vs vehicle-treated comparison. Pathways displayed are significantly differently regulated (adjusted p-value ≤ 0.05 by Fisher’s test). The length of bars reflects the number of genes in the pathway whose phosphostate is differentially regulated; bars are shaded by adjusted p-value. Highlighted pathways are related to established LRRK2 functions. **D.** Diagram of genetic and pharmacological conditions for groups analyzed in E. Created with BioRender.com. **E.** Heatmap of effect size (Log2FC) of either haloperidol treatment or LRRK2-GS for all differentially abundant phosphopeptides in the WT haloperidol-treated vs vehicle-treated comparison (|Log2FC| > 2 and unadjusted p-value ≤ 0.05 by multiple unpaired t-tests). Each bar represents a phosphopeptide. Color reflects Log2FC difference from WT saline condition. **F.** Correlation plot for E, comparing GS vehicle/WT vehicle to WT haloperidol-vehicle effect size. All detected phosphopeptides were mapped, and phosphopeptides significantly altered between WT haloperidol vs. vehicle are highlighted. Only highlighted values are used for correlation calculation. Blue line represents the line of best fit.

The timeline of effects of LRRK2 inhibition on the cataleptic response (7 days of treatment) correlates with a time when synaptic adaptations in the iSPNs have started to occur. Moreover, although the cataleptic tolerance progressively increased from 7 to 14 days in haloperidol-treated WT mice (p=0.0226, Tukey’s multiple comparisons test comparisons test after 2-way ANOVA), no difference in haloperidol induced catalepsy was observed in the MLi-2-treated mice between 7 and 14 days (Fig. 2B), suggesting that LRRK2 kinase activity interferes with the well-orchestrated changes in iSPNs pathways after haloperidol treatment. Thus, we examined the haloperidol-induced dendritic spine density patterns in iSPNs after 7 days of treatment. We identified iSPNs by injecting a Cre-dependent adeno-associated virus (AAV) expressing eGFP (AAV5/DIO-EGFP) in the striatum of *Adora2a^Cre^* mice (Fig. 2C, Suppl. Fig. 4). Consistent with previous reports^16^, we found that haloperidol decreased dendritic spines density on iSPNs, compared to the vehicle control (p<0.0001, Tukey’s multiple comparisons test after one way ANOVA, n=14-20 iSPNs), (Fig. 2C, D). Co-administration of MLi-2 and haloperidol increased dendritic spine density in iSPNs, compared to iSPNs after haloperidol alone (p=0.0042, Tukey’s multiple comparisons test after one-way ANOVA, n=14-20 iSPNs), (Fig. 2C, D). Thus, LRRK2 kinase activity is involved in the synaptic changes in iSPNs circuits following haloperidol, consistent with a well-described role for LRRK2 in dendritic spine morphogenesis through actin cytoskeleton regulation.

To better understand LRRK2 effects on intrinsic iSPNs adaptations, *Drd2-eGFP* BAC reporter mice were injected with haloperidol, or haloperidol + MLi-2, or vehicle controls for 14 days. Then, we performed whole-cell current-clamp recordings on identified iSPNs in ex vivo striatal slices (Fig. 2E). Our findings are in line with a previous study^16^ that showed a decrease in iSPNs excitability due to haloperidol (treatment group by current interaction, p=0.0001, 2-way ANOVA with Sidak multiple comparisons, n=4-6 animals, n=14-24 iSPNs). Importantly, co-administration of MLi-2 with haloperidol restored the firing of iSPNs to wild-type levels (p=0.9506, 2-way ANOVA with multiple comparisons, n=4-6 animals, 14-29 iSPNs) (Figs 2E-F). Our results indicate that LRRK2 kinase activity is involved in synaptic and intrinsic changes in iSPN circuits after haloperidol treatment.

### Haloperidol administration mimics the phosphoproteomic landscape of the LRRK2 G2019S mutation in the striatum

Previous studies have shown that the phosphorylation of ribosomal protein S6 (p-rpS6) at S235/236 residues increases in iSPNs in response to haloperidol activation^54,55^. This phosphorylation is cAMP/PKA dependent, which is the canonical pathway inhibited by D2R antagonism. We therefore explored whether LRRK2 inhibition interferes with this haloperidol-mediated phosphorylation event.

Mice expressing the eGFP under the *Drd2* promoter were treated with haloperidol, MLi-2, haloperidol plus MLi-2, or vehicle controls for 7 days (Suppl. Fig. 5A), and double immunofluorescence experiments in striatal sections using antibodies against the p-rpS6 and GFP (for iSPNs) were performed. As expected, haloperidol increased the percentage of pS6-Ser235/236 positive iSPNs in striatal sections (p<0.0001, Tukey’s multiple comparisons test after one-way ANOVA, n=21-22 sections). In mice treated with both MLi-2 and haloperidol, the number of p-rpS6 GFP+ neurons was significantly lower than in mice treated only with haloperidol, and similar to vehicle controls (p=0.2311, Tukey’s multiple comparisons test after one-way ANOVA, n=18-22 sections) and (p=0.0019, Tukey’s multiple comparisons test after one-way ANOVA, n=18-21 sections), (Suppl. Fig. 5B). Unexpectedly, when MLi-2 was administered alone, p-rpS6 GFP+ neurons increased compared to the control group (p=0.0476, Tukey’s multiple comparisons test after one-way ANOVA, n=18-22 sections). These results indicate that inhibiting LRRK2 kinase activity blunts the intracellular response to haloperidol, which correlates with the observed improvement in haloperidol-induced behavioral responses in the presence of LRRK2 kinase inhibitors.

Since LRRK2 kinase inhibition reduces the intracellular response to haloperidol, we predicted that hyperactive mutant LRRK2 should mimic haloperidol-induced molecular alterations in WT mice. To test this hypothesis, we conducted unbiased proteomic characterization in the striatum, using mice expressing the G2019S (GS) mutation of LRRK2, which increases its kinase activity. We acutely (1 hour) administered haloperidol or vehicle to WT and GS KI mice, extracting striatal samples for quantitative total and phosphoproteomic analyses (Fig. 3A). Our proteomic analysis detected 3,997 proteins, while the phosphoproteomic analysis detected 9,399 phosphopeptides mapping to 3,010 unique proteins. For phosphoproteomics, the log2 intensity level of each phospho-peptide was normalized over the log2 intensity level of the total corresponding protein. We first compared haloperidol and vehicle-treated WT mice, and we found 252 phosphopeptides that were differentially regulated (p-value < 0.05, & |Log2 fold change (FC)| > 2) (Fig. 3B). Gene set enrichment analysis (GSEA) of the KEGG pathway set was conducted on genes containing at least one significantly altered phosphopeptide. Several KEGG pathways were found to be differentially regulated, including glutamatergic synapses, cAMP signaling, and actin dynamics (Fig.3C). These processes have been previously linked to the effects of haloperidol on the striatum^54,56–59^. Several of the observed differentially altered pathways, such as cAMP/PKA signaling and the regulation of actin cytoskeleton, represent established roles for LRRK2 function^26,20,61^. Specifically, LRRK2 involvement in regulating actin cytoskeleton, which underlies dendritic spine morphogenesis, has been well-established^62–66^. Additionally, we have previously shown that LRRK2 directs synaptogenesis via a PKA-mediated mechanism^20^. Further ontology analyses of significant phosphopeptides supported the enrichment of cytoskeletal and postsynaptic organization-associated pathways (Suppl. Fig. 6A).

These findings suggest that haloperidol-induced motor changes involve modulation of LRRK2 kinase activity. Further supporting this model, phosphopeptides found to be differentially abundant in haloperidol-treated vs. vehicle-treated WT mice show changes similar in direction and magnitude as phosphopeptides differentially abundant in vehicle-treated LRRK2 GS vs. WT mice (Fig. 3D-E), with a strong correlation of effect size (Fig. 3F). These results indicate that haloperidol treatment results in similar phosphoregulation patterns in the striatum compared to those caused by hyperactive LRRK2. While the phosphoproteomic changes resulting from haloperidol treatment were almost wholly reflected in the changes as a result of hyperactive LRRK2, there was a large population of phosphopeptides that were significantly differentially regulated in the GS mice but unchanged as a result of haloperidol treatment (Suppl. Fig. 6B). This suggests that chronic hyperactivity of LRRK2 in GS mice results in more profound phosphorylation restructuring and reflects broader functions related to the LRRK2 kinase as compared to the changes observed upon transient administration of antipsychotic D2R antagonists.

In contrast to the phosphoproteomic data, analysis of changes in total protein levels between wild-type mice treated with haloperidol and vehicle uncovered very few differentially expressed proteins (Suppl. Fig. 6C). These significantly altered proteins did not show similar changes in expression in vehicle-treated LRRK2-GS mice compared to WT (Suppl. Fig. 6D). Finally, a moderate effect of haloperidol on the phosphoproteome was observed when comparing vehicle-treated and haloperidol-treated LRRK2-GS mice (Suppl. Fig. 6E), consistent with the idea that haloperidol mimics LRRK2 hyperactivity. These results demonstrate that haloperidol administration primarily impacts striatal phosphostate due to changes in intracellular signaling, with LRRK2 kinase activity playing a key role in this state transition.

### LRRK2 regulates haloperidol mediated upregulation of immediate early genes

Antipsychotics are known to induce the expression of several immediate early genes (IEGs) via D2R antagonism, which then leads to changes in critical signaling pathways in iSPNs^67,68^. Although the exact intracellular pathways leading to this upregulation are unclear, it is known that D2R blockade by haloperidol increases the activity of PKA (Fig. 4A). This then causes phosphorylation of CREB, allowing it to bind to various IEG promoters to induce transcription^54,69, 55,70^.

**Figure 4.**
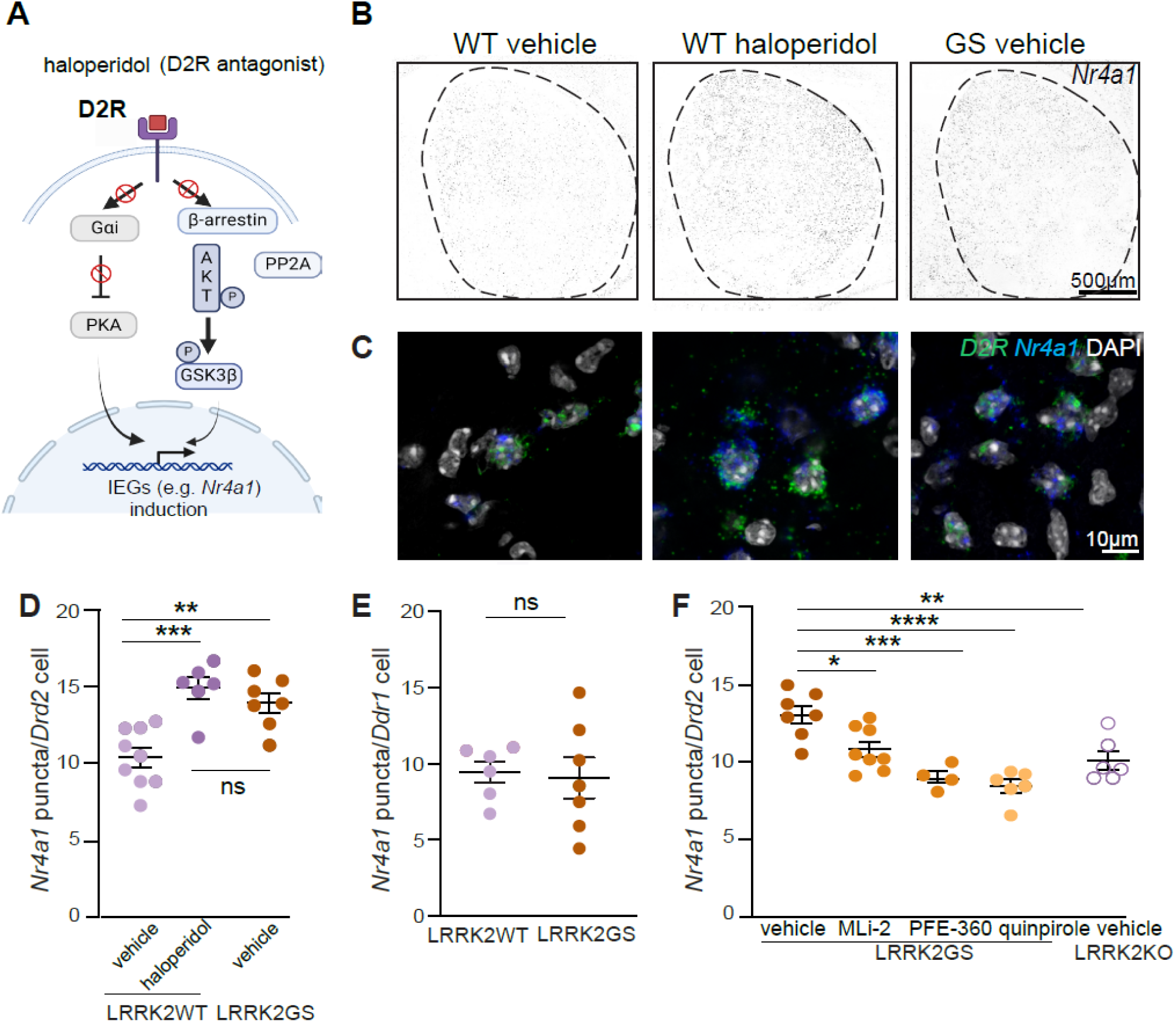
LRRK2 regulates the D2R mediated increase of *Nr4a1* in indirect pathway SPNs. **A.** Signaling cascades leading to IEGs upregulation in iSPNs. D2R antagonist (haloperidol) releases the inhibition of cAMP and activates PKA, leading to upregulation of IEGs (canonical pathway). D2R blockade prevents the reduction of AKT activity, increasing GSK3β phosphorylation/inactivation. Inhibition of GSK3β elicits IEG expression. Created with BioRender.com. **B.** Example confocal images of *Nr4a1* gene expression in the striatum of LRRK2-WT mice treated with haloperidol (1 mg/kg, for 1 hour) or vehicle and LRRK2-GS mice treated with vehicle. Scale bar=500 μm. **C.** Same as B, but high-magnification images depicting *Nr4a1* expression. Scale bar=10 μm **D.** Comparison of mean puncta/cell of *Nra41* in *Drd2*-positive nuclei of LRRK2-WT mice treated with haloperidol or vehicle and LRRK2-GS mice treated with vehicle. Each dot represents the average number of *Nra41* positive cells from one striatal section, n=6-9 sections/3-4 mice. The number of *Nr4a1+*/*Drd2+*positive cells across WT and GS mice does not differ (Tukey’s multiple comparison tests after two-way ANOVA). **E.** Comparison of mean puncta/cell of *Nra41* in *Drd1*-positive nuclei of LRRK2-WT mice and LRRK2-GS mice treated with vehicle. Each dot represents the average number of Nra41 positive cells from one striatal section, n=6-7 sections/3 mice. The number of *Nr4a1+/Drd1+* positive cells across WT and GS mice does not differ (unpaired t-test). **F.** Quantification of *Nra41*-positive *Drd2* nuclei in LRRK2-GS mice administered vehicle, MLi2 (10 mg/kg), PFE-360 (5 mg/kg), D2R agonist quinpirole (1 mg/kg), as well as LRRK2-KO mice treated with vehicle for 2 hours. N=4-8 sections/2-4 mice Data are represented as mean±SEM. Asterisks in F reflect statistical significance for Tukey post-hoc comparisons after one-way ANOVA. * p<0.05, *p < 0.01, **p< 0.001 ****p < 0.0001.

Nuclear receptor subfamily 4 group A member 1 (Nr4a1) is a transcription factor and IEG that is strongly upregulated in the iSPNs of the dorsolateral striatum after haloperidol treatment^71,72^. Since the dorsolateral striatum is crucial for motor control, the upregulation of *Nr4a1* after haloperidol administration may be associated with subsequent motor side effects. Indeed, genetic loss of *Nr4a1* leads to the complete blockade of haloperidol-induced catalepsy^73^.

Therefore, we explored whether LRRK2 kinase is part of the intracellular signaling cascade downstream of D2R that leads to haloperidol mediated *Nr4a1* induction. To define the impact of LRRK2 kinase activity on the expression pattern of *Nr4a1* mRNA in the iSPNs, we used multicolored single-molecule fluorescence in situ hybridization. In addition to comparing WT to LRRK2-GS iSPNs, we compared WT mice to those treated with haloperidol (Fig. 4B, C). Consistent with previous work^72^, we found an upregulation of *Nr4a1* in the iSPNs of WT mice upon haloperidol administration, defined by the number of *Nr4a1* puncta per iSPN (p=0.0002, Tukey’s multiple comparisons test after one-way ANOVA, n=6-9 sections). Similarly, we observed increased *Nr4a1+* puncta in GS iSPNs compared to the WT controls (p=0.0018, Tukey’s multiple comparisons test after one-way ANOVA, n=7-9 sections) (Fig. 4D). In contrast, we found no difference in the number of *Nr4a1* puncta in the dopamine D1 receptor (D1R)-expressing SPNs in LRRK2-GS mice (p=0.8193, unpaired t-test, n=6-7 sections, Fig.4E), indicating a pathway-specific effect. Two different LRRK2 kinase inhibitors, MLi-2 and PFE-360, were separately delivered orally to GS KI mice (2 hours). This restored (decreased) the *Nr4a1* expression in LRRK2-GS mice, compared to vehicle controls (p=0.0237 MLI-2, p=0.0003 PFE-360, Tukey’s multiple comparisons test after one-way ANOVA n=4-8 sections, Fig. 4F). Moreover, the number of *Nr4a1+* puncta in iSPNs of vehicle-treated LRRK2-KO mice was decreased, compared to LRRK2-GS vehicle-treated mice (p=0.0019, Tukey’s multiple comparisons test after one-way ANOVA n=6-7 sections, Fig. 4F). To assess whether decreased D2R signaling mediates the enhanced *Nr4a1* mRNA levels in LRRK2-GS iSPNs, we systematically administered the D2R agonist quinpirole. D2R stimulation decreased the levels of *Nr4a1* in LRRK2-GS iSPNs (p<0.0001, Tukey’s multiple comparisons test after one-way ANOVA, n=6-7 sections) compared to vehicle controls (Fig. 4F). These results demonstrate a LRRK2 kinase-mediated and D2R signaling-dependent upregulation of *Nr4a1* in the LRRK2-GS iSPNs.

If LRRK2 regulates IEG expression through modulation of the D2R signaling, the effects of inceased LRRK2 kinase activity should be generalizable for any IEG that responds to D2R signaling, even across IEGs that encode proteins with diverse functions. In addition to the expression levels of *Nr4a1*, which encodes a transcription factor, we examined the number of *Arc* (encoding a dendritic spine protein) positive puncta in iSPNs of LRRK2-GS mice and WT mice treated with haloperidol (Suppl. Fig. 7A). Using in situ hybridization, we found an increase in *Arc*-positive puncta in the iSPNs of haloperidol-treated compared to vehicle-treated WT mice (p=0.0056, Tukey’s multiple comparisons test after one-way ANOVA n=4-7 sections) (Suppl. Fig. 7B). We also observed higher *Arc+* puncta in iSPNs from LRRK2-GS, compared to WT (p=0.0002, Tukey’s multiple comparisons test after one-way ANOVA as noted in Suppl. Fig. 7B, n=4-7 sections). To determine whether this increase is LRRK2 kinase-dependent, we treated LRRK2-GS mice with the kinase inhibitor MLi-2, which decreased the *Arc* RNA levels, compared to vehicle-treated LRRK2-GS iSPNs (p=0.0101, Tukey’s multiple comparisons test after one-way ANOVA, n=6-7 sections), (Suppl. Fig. 7C) Moreover, we found that LRRK2-KO iSPNs had less *Arc*+ puncta/cell than LRRK2-GS iSPNs (p=0.0251, Tukey’s multiple comparisons test after one-way ANOVA, n=6-7 sections), (Suppl. Fig. 7C). Together, our data show that–similar to haloperidol treatment–genetically increased LRRK2 kinase activity results in a D2R-dependent upregulation of several IEGs, which reflects a broad effect of LRRK2 on signaling modalities downstream of D2R.

## Discussion

This study shows that LRRK2 kinase activity is involved in D2R-mediated motor behaviors and their modulation by antipsychotic D2R antagonism. We employed both pharmacological (potent and specific LRRK2 kinase inhibitors) and genetic (LRRK2-KO and LRRK2-GS KI mice) models to modulate levels of LRRK2 kinase activity. We demonstrated that pharmacological or genetic inhibition of LRRK2 kinase activity attenuates the behavioral, electrophysiological, and synaptic effects of haloperidol administration. Furthermore, we characterized the proteomic and phosphoproteomic changes induced by haloperidol treatment in the striatum and found the impact of increased LRRK2 kinase activity mirrored them. In addition, increased LRRK2 kinase activity contributed to intracellular pathway responses of iSPNs to haloperidol treatment. Our study suggests that LRRK2 kinase activity is involved in critical D2R-mediated signaling events, broadly relevant to PD pathophysiology. Furthermore, our results highlight the potential of LRRK2 inhibitors to ameliorate extrapyramidal motor side effects of D2-antagonistic antipsychotics.

### Converging effects of haloperidol and LRRK2 kinase activity on changes in striatal indirect pathway Behavioral and anatomical insights

We observed that catalepsy and reduced locomotion induced by haloperidol treatment were attenuated in the presence of pharmacological or genetic LRRK2 inhibition (Fig. 1). Although we cannot rule out the potential effects of LRRK2 on other D2R-^55^ or D1R-expressing^74^ striatal subpopulations, our findings show that knockout of the long isoform of D2R^15^, which is predominantly expressed in iSPNs, decreases substantially the cataleptic response to haloperidol. This demonstrates a critical contribution of D2R signaling in iSPNs to the cataleptic response.

Haloperidol has been shown to induce synaptic changes in the indirect striatal pathway, including a decrease in the density of dendritic spines^16^. Our data demonstrate that this decrease is partially restored by inhibiting LRRK2 (Fig. 2). LRRK2 plays a crucial role in dendritic spine morphogenesis, through PKA-dependent regulation of the actin cytoskeleton^20^. Actin-dependent remodeling of dendritic spines as a result of haloperidol administration is supported by our phosphoproteomic data, which showed significant alterations in the phosphostate of actin cytoskeletal-related proteins with important roles in dendritic spine formation, such as drebrin, cofilin, and neurabin after haloperidol treatment. Altogether, our results suggest that LRRK2 and haloperidol converge on regulating actin remodeling as a functional downstream effect of D2R antagonism.

### Molecular mechanisms

Unbiased phosphoproteomics in striatal extracts from haloperidol-treated mice revealed changes in phosphoregulation of cAMP/PKA signaling-linked pathways, which are the canonical pathways downstream of D2R activation, as well as changes in the phosphorylation of proteins linked to actin cytoskeleton dynamics and glutamatergic synapses (Fig. 3). These findings align with structural and functional changes previously reported to result from haloperidol treatment in corticostriatal glutamatergic synapses^16,52^. Our findings showed a larger magnitude of phosphoproteomic changes compared to a largely stable proteome. This stands in contrast to prior findings reporting substantial proteomic changes as a result of haloperidol treatment^56^. These differences are likely due to the temporal dynamics of haloperidol administration in each study. We chose acute treatment to capture the initial signaling dynamics following D2R antagonism, as this is where phosphoregulation by LRRK2 kinase is expected to be involved. In contrast, Santa and colleagues employed a 30-day chronic haloperidol treatment paradigm. Our combined findings suggest that phosphoproteomic alterations precede the subsequent reshaping of the proteomic landscape that occurs with continued haloperidol treatment.

Although we cannot currently understand how LRRK2 kinase activity phenocopies the effects of haloperidol treatment, our data point to convergent alterations in signaling downstream of D2Rs. LRRK2 activity has been shown to impact the intracellular trafficking of D2Rs^75^, which in this manner may mimic the downstream intracellular cascades that are affected by the direct action/blockade of antipsychotics on D2Rs. In addition, we find that expression of the hyperactive LRRK2-GS mutation mimics the increase in IEG mRNAs such as *Nr4a1* and *Arc* in iSPNs (Fig. 4). This may directly contribute to the cataleptic tolerance upon LRRK2 kinase inhibition since *Nr41a1* knockout mice have been shown to have a blunted cataleptic response^73^. The selective induction of IEGs in the GS mice is a striking example of a pathway-specific signaling effect, in line with our previous anatomical, physiological, and behavioral studies that support pathway-specific effects of LRRK2^26,23^ .

### Clinical implications

Our study shows that inhibiting LRRK2 kinase activity decreases locomotor deficits as a result of D2R antagonism via haloperidol treatment. This finding has two significant clinical implications. First, under conditions of increased kinase activity, such as in individuals with GS mutations, stimulation of D2R signaling may act to reverse the negative pathological effects of hyperactive LRRK2. Our data show that quinpirole, a D2R agonist, attenuates the increased expression of IEGs in GS iSPNs (Fig. 4). This is consistent with other studies showing that stimulation of D2R signaling is beneficial in the context of enhanced LRRK2 kinase activity^27,28^ with significant implications on the clinical use of DA agonists in PD^76^. Second, LRRK2 kinase inhibition may ameliorate the serious Parkinsonian-like motor phenotypes induced by haloperidol. Our data suggest a framework by which LRRK2 kinase inhibition^77^ or anti-sense oligonucleotides to LRRK2^78^, both of which are currently in clinical development, could be used as adjuncts to antipsychotics. This is a viable therapeutic opportunity as LRRK2 loss of function variants in humans are not associated with disease state or dysfunction^79^. It remains to be determined whether LRRK2 kinase activity is relevant to the therapeutic benefits of haloperidol. The delayed clinical effect of haloperidol (weeks or months) suggests that it ameliorates psychotic symptoms through mechanisms beyond the acute blockade of D2R, potentially involving network-level adaptations^80,81^. Investigating the effects of LRRK2 inhibition on the complex and less well-understood effect of haloperidol on the positive symptoms of schizophrenia would be valuable in future studies. Our work is an important step in understanding the neurobiology of dopamine-related disorders and their connection with motor symptoms, paving the way for more effective therapeutics for these neurological conditions.

## Acknowledgments

This work was supported by NIH R01 NS097901 (L.P.), R01NS107539 (Y.K.) and 2021 OneMind Nick LeDeit Rising Star Research Award (Y.K.). This research was funded in whole or in part by Aligning Science Across Parkinson’s [ASAP-020600] through the Michael J. Fox Foundation for Parkinson’s Research (MJFF) (L.P.). For the purpose of open access, the author has applied a CC BY public copyright license to all Author Accepted Manuscripts arising from this submission. Research reported in this publication was supported by the National Institute of General Medical Sciences of the National Institutes of Health under Award Number T32GM105538 (M.M.). The content is solely the responsibility of the authors and does not necessarily represent the official views of the National Institutes of Health.

## Authors contributions

Conceptualization, C.C., L.P.; scientific input, D.A., E.G., S.H., and Y.G.; methodology, C.C., M.M., N.S., V.P., G. T., C.M.; molecular and anatomical methods C.C; electrophysiological measurements, M.M. and Y.K.; behavioral tests, C.C., V.P., G.T., data curation, formal analysis, C.M.; formal analysis, N.S., writing first draft, C.C., and L.P.; writing , review and editing, all authors; funding acquisition, L.P. Y.K.; supervision, L.P.

## Data sharing

The mass spectrometry data will be deposited to the ProteomeXchange Consortium via the PRIDE partner repository; primary data that is presented in this study will be submitted to Zenodo; protocols described in this paper will be deposited to protocols.io before formal manuscript acceptance.

## Resources Table

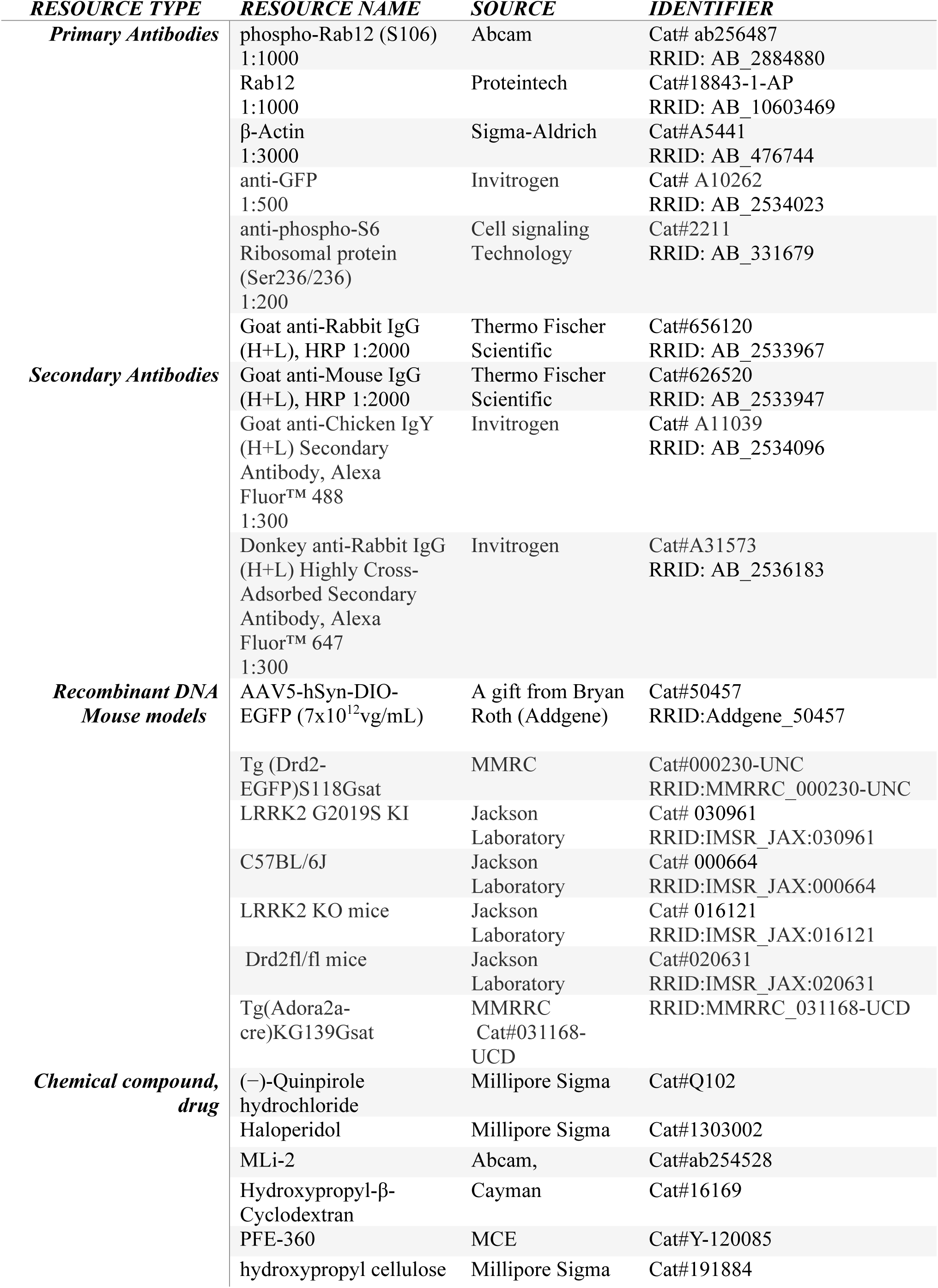
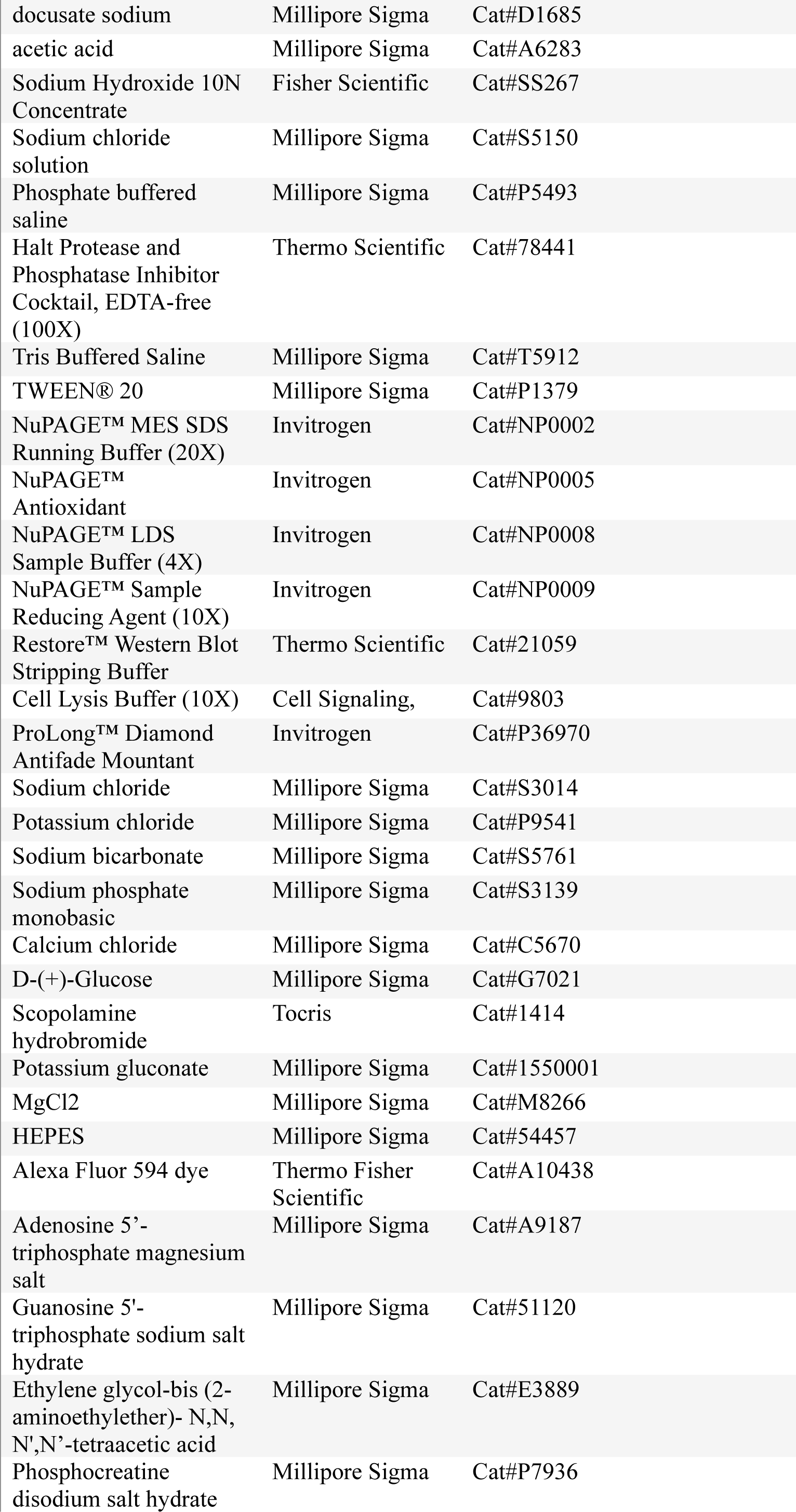
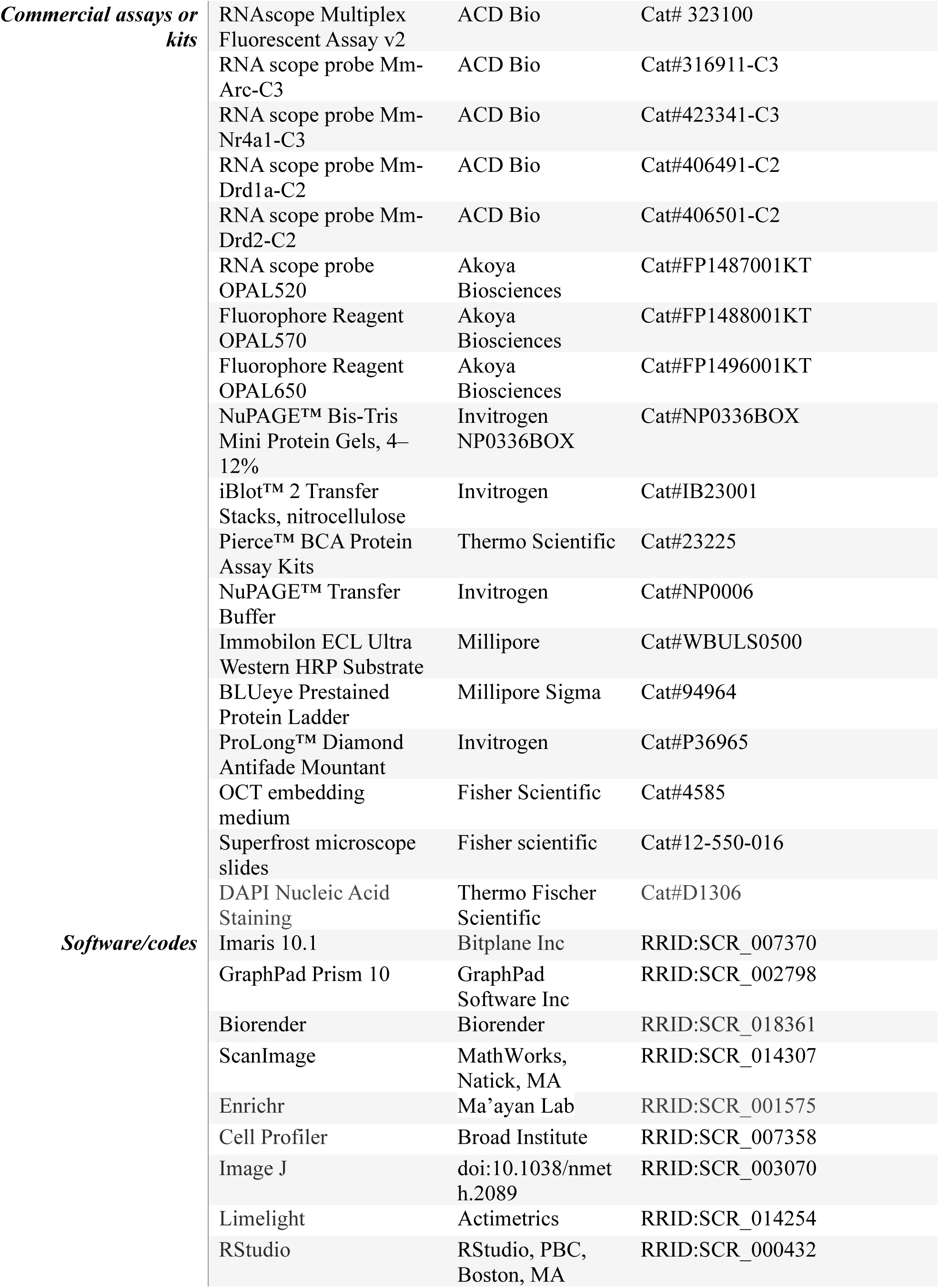

## Materials and methods

### 1. Animals

All experiments were in compliance with National Institutes of Health guidelines and were reviewed by the Northwestern Animal Care and Use Committee. P28-P35 male and female mice were used in all experiments. The effect of sex for key behavioral experiments is summarized in Suppl. Fig. 2. Mice were group-housed on a standard 12/12 hr light/dark cycle with standard feeding. Littermates were randomly assigned to the experimental procedures. C57BL/6 and heterozygous KI mice expressing the LRRK2-GS pathogenic mutation^33^ referred to as LRRK2-GS, as well as homozygous LRRK2-KO mice^65^ were from Jackson Laboratory. BAC transgenic mice in which EGFP expression is driven by dopamine receptor D2 *Drd2* regulatory elements (GENSAT)^82^ referred to as *Drd2-eGFP* allow the visual identification of these SPNs for electrophysiological (Fig. 2) and immunofluorescence studies (Suppl. Fig.5). These mice crossed with LRRK2-GS KI mice were used in heterozygosity in all experiments. GENSAT generated^82^*Adora2a* (KG139) BAC mice which express Cre recombinase under the control of the Adenosine A2a receptor promoter, *(Adora2a^Cre^ )* were crossed to a mouse line with *loxP* sites flanking exon dopamine receptor D2 (*Drd2*) gene, referred to as *Drd2^fl/fl^* mice^83^ and used in behavioral tests (Fig.1) Conditional expression of fluorescent proteins in *Adora2a^Cre^ (iSPNs)*-containing neurons for dendritic spines analysis (Fig. 2) was achieved using adeno-associated viruses encoding a double-floxed inverted open reading frame (DIO) of eGFP referred to as AAV-DIO-EGFP. The Cre+ mice used were heterozygous. All mice were back crossed several generations and maintained on a C57BL/6J background. Standard genotyping primers are available on the Jackson Lab or MMRC website. All animals were genotyped according to the Jackson Laboratory or MMRRC strain-specific primers and protocols Details about the mouse strains are in the resources table.

### 2. Drugs and treatment dosing

Haloperidol (1mg/Kg) was dissolved in 0.05% acetic acid in saline (pH adjusted to 6.0 with NaOH), quinpirole (1 mg/kg) was dissolved in saline. Both drugs and corresponding vehicle controls were administered intraperitoneally (i.p). Mli-2 (10mg/Kg) was dissolved in 40% (w/v) Hydroxypropyl-β-Cyclodextran, and PFE-360 (5mg/Kg) in 1.25% hydroxypropyl cellulose + 0.5% docusate sodium. Mli-2, PFE360, and their vehicle controls were administered via oral gavage. Drugs and all reagents used are detailed in the Resources Table, along with identification numbers when applicable.

#### Acute treatments

Mice were given a dose of MLi-2 or PFE-360 or vehicles via oral gavage 30 min before i.p. haloperidol injections or its vehicle for an hour. At this time point, the mice were euthanized for subsequent behavioral (Fig.1), immunoblotting (Suppl. Fig.1), mass spectrometry (haloperidol for 1 hour) (Fig. 3) or in situ hybridization (Fig. 4) studies. Quinpirole was administered for 1 hour (Fig. 4).

#### 7- and 14-day treatments

Mice were administered haloperidol following MLi-2 or PFE-360 as in the acute paradigm once a day for 7 or 14 days. They were euthanized for behavioral (Suppl. Fig 1), immunoblotting (Suppl. Fig. 1), and immunofluorescence (Suppl. Fig. 5) studies 1 hour after the last injection. The indicated experimental procedures for the haloperidol-mediated synaptic and intrinsic adaptations (Fig. 2) and the rotarod test in Suppl. Fig. 3 were performed 24 hours after the last haloperidol injection.

### 3. Behavioral Analyses

#### Bar test

Mice were gently removed from their home cage, and their forepaws were placed over a horizontal bar, 0.2 cm in diameter, fixed at a height of 4 cm above the working surface. The time a mouse remained immobile was measured in 3 trials, with the cut-off at 120 sec^15^. The average of 3 trials was used for further statistical testing.

#### Open field

Mice were put in a 56 ×56 cm open-field arenas in noise-canceling boxes, illuminated by dim red lights. The 20-minute-long session started when mice were placed in the center of the arena. Locomotor activity was analyzed by the LimeLight 5 (Actimetrics) software and reported as distance traveled.

#### Rotarod

Motor learning was assessed with an accelerating rotarod. The task was done using a rotarod apparatus (Panlab) equipped with a mouse rod (3 cm diameter) and set to 4–40 rpm acceleration over 300 s. The task consisted of daily sessions (five trials per session; intertrial-interval = 15 s, max trial duration = 300 s). Following a 24-hour break, mice were tested for 8 sessions.

### 4. Stereotaxic Surgeries

#### Neonatal

For dendritic spine analysis in Fig. 2, GS mice were crossed with *Adora2a^Cre^* mice. Intracranial injections were performed as previously described ^84–86, 26^. P4-5-day-old pups were cryoanesthetized and received ketoprofen for analgesia. They were placed on a cooling pad on a stereotaxic frame. 200 nl of the AAV-DIO-EGFP virus (7×10¹² vg/mL) were delivered into the dorsal striatum at a rate of 100 nl min^-1^ using Micro-4 controller (WPI). In order to ensure the dorsal striatum was targeted, the needle was placed 1 mm anterior to the midpoint between the ear and eye, 1.5 mm from the midline, and 1.8 mm ventral to the brain surface. Pups’ age and size determined slight adjustments in the coordinates. After the injection, the pups were warmed on a heating pad before returning to their home cages.

### 5. Confocal microscopy and dendritic spine analysis

Confocal images of fixed 80-um-thick brain sections of P30 pups injected with the AAV-DIO-EGFP (Fig, 2) were obtained with the Nikon A1R microscope. Fluorescence projection images of dendrites and the corresponding dendritic spines were acquired with a 60x oil immersion objective (NA = 1.4) at 0.1 um intervals with 1,024 x 1,024 pixel resolution. For each group, 2–4 segments per neuron 5-9 neurons from 2-3 animals were used to generate z-stacks. Segments from secondary and tertiary dendrites without overlap with other neurons or discontinuities were chosen for analysis. Dendritic spine density was assessed using Imaris 10.1 software (Bitplane, Concord, USA). Images of dendritic segments were traced using autopath mode of the filament tracer at default settings. The following settings were used for spine detection: thinnest diameter of seed points at 0.45 μm; maximum length at 2.5 μm; no branched spines allowed; and seed point threshold at auto.

### 6. Acute Slice Preparation

*Drd2-eGFP* mice were treated with either haloperidol or haloperidol plus LRRK2 inhibitor (MLi-2) as described above for 14 days. On day 15 or 16, mice were deeply anesthetized on isoflurane (3%) prior to perfusion with ice-cold aCSF containing in (mM): 127 NaCl, 2.5 KCl, 1.25 NaH_2_PO_4_, 25 NaHCO_3_, 20 glucose, 2 CaCl_2_, and 1 MgCl_2_. Brains were dissected and sectioned on a Leica vibratome (VT1000S) to 300 um thickness. Slices were transferred to a holding chamber and incubated at 34°C for 20 minutes prior to recordings.

### 7. Current clamp recordings

Slices were placed in the recording chamber at RT and continually perfused with oxygenated aCSF with 1mM scopolamine to block cholinergic interneuron firing. The dorsal striatum was located under a 10x air objective, then switched to a 60x water immersion objective was used to target neurons for recording (Olympus, Tokyo, Japan). Slices were visualized using DIC and a QIClick microscope camera (QImaging, Surrey, Canada). GFP-positive cells were selected using a CoolLED pe4000 system (CoolLED Ltd., Andover, UK). Patch electrodes were pulled from borosilicate capillary glass to a resistance of 2.5–6 MΩ. Electrodes were filled with an intracellular solution comprised of (in mM) 135 K-gluconate, 4 KCl, 10 HEPES, 10 Na-phosphocreatine, 4 MgATP, 0.4 Na_2_GTP, 1 EGTA, 20 µM Alexa 594 (pH 7.28, 298-305 mOsm/L). After break-in, cells were held for ∼5-10 minutes prior to current clamp recordings.

Recordings were acquired using a Multiclamp 700B amplifier (Axon Instruments, Union City, CA), digitized at a rate of 20 kHz, and filtered at a rate of 3-4 kHz. Five second-long sweeps were acquired using a version of the MATLAB-based (MathWorks, Natick, MA) acquisition suite, ScanImage^87^. A series of 500 ms-long current steps from -50 to 300 pA were pseudorandomized. Current steps occurred one second after the start of the sweep, with a 20 ms-long, -20 pA current step added at the end of the sweep to test input resistance throughout the recording. Cells with input resistance that changed >10% over the recording were discarded. Sweeps occurred with an inter-sweep interval of 10-20 seconds. A minimum of five sweeps for each current step magnitude was obtained.

### 8. Current clamp recording analysis

All acquisition and data analysis were done blind to treatment conditions. Average firing rate at each current step was determined using custom built MatLab code. All values are reported as Hz. Firing rate values were then exported to GraphPad Prism (GraphPad, LaJolla, CA) for data visualization and statistics. Two-way ANOVA with Geisser-Greenhouse correction for symmetry was used to determine significance of treatment by current injection interaction, with post hoc multiple comparisons with Šídák correction across current injection levels.

### 9. Western blot analysis

After receiving the drug treatments as described in the relevant sectionh striatal tissues were extracted and homogenized in 1x cell lysis buffer (Cell Signaling Technologies) supplemented with Halt protease and phosphatase inhibitor cocktail (Thermo Fisher Scientific) using pellet pestles (30 seconds). Protein concentration was determined by BCA Protein Assays (Thermo Scientific) 30 μg of total lysate were separated by 4–12% NuPage Bis-Tris PAGE (Invitrogen) and transferred to membranes using the iBlot nitrocellulose membrane Blotting system (Invitrogen) by following manufacture protocol. The membranes were incubated with primary antibodies specific for phospho-Rab12 (1:1000, Abcam), total Rab12 (1:1000 Proteintech and β-actin (1:3000, Sigma) at 4C overnight. The membranes following probing with secondary anti-mouse or and anti-rabbit antibodies (1:2000, Thermo Scientific) for 1 hour at room temperature were incubated with Immobilon ECL Ultra Western HRP Substrate (Millipore) for 3 min prior to image acquisition. Chemiluminescent blots were imaged with iBright CL1000 imaging system (Thermo Fisher Scientific). Chemicals and antibodies are detailed in the resource table.

### 10. Striatal sections immunofluorescence and analysis

*Drd2-eGFP* mice crossed with LRRK2-GS KI and control littermates were treated as indicated and perfused with 50 ml of PBS and followed with 50 ml 4% paraformaldehyde in PBS 1 h after the last vehicle/haloperidol administration. Brains were dehydrated with 30% sucrose in PBS for 48hrs and cut coronally (30 μm) by cryostat (Leica Biosystems,CM305). Slices were collected in PBS + 0.1% sodium azide and stored at 4°C for immunohistochemistry. Striatal sections were incubated with 5% goat serum in 0.2% Triton X-100 for 2 hrs. After, the sections were incubated in the same solution overnight at 4 °C with the primary antibodies anti-GFP (1:1000, Invitrogen) and anti-phospho-S6 Ribosomal protein (Ser236/236) (1:300, Cell Signaling Technology). Then, sections were washed with PBS and incubated with the secondary antibodies Alexa Fluor™ 488 and Alexa Fluor™ 647 (both 1:600, Invitrogen) for 3 hrs. All the slices were washed with PBS and mounted on ProLong™ Diamond Antifade Mountant. Detailed catalog and identification numbers are shown in the resource table.

Confocal images were obtained with a Nikon A1R microscope. Fluorescence images were acquired with a 20x objective at 1,024 x1,024 pixel resolution. Stitched images of the whole striatum were automatically acquired with Nikon Element. GFP and phospho-S6 Ribosomal protein signal were measured using Imaris 10.1 software (Bitplane, Concord, USA). Surface rendering function was used to segment Drd2-eGFP cells. Background subtraction was enabled, the diameter of the largest sphere was set at 15 μm and automatically thresholded, and smooth surface was set to 1.23 μm. Segments were filtered with an area ranging between 100 μm^2^ and 6,000 μm^2^. Mean intensity of phospho-S6 Ribosomal protein within Drd2-eGFP surface above 2 times of average phospho-S6 Ribosomal protein channel mean intensity was considered positive.

### 11. LC-MS/MS Analysis

The tissue samples were processed by Tymora Analytical Operations (West Lafayette, IN). For lysis, 200 µL of phase-transfer surfactant lysis buffer (PTS, containing 12 mM sodium deoxycholate, 12 mM sodium lauroyl sarcosinate, 10 mM TCEP, 40 mM CAA), supplemented with phosphatase inhibitor cocktail 3 (Millipore-Sigma) was added to each of the tissue samples, pulse-sonicated several times with a sonicator probe to lyse the tissues. The samples were incubated for 10 min at 95°C, pulse-sonicated several times again with a sonicator probe, and incubated again for 5 min at 95°C. The lysed samples were then centrifuged at 16,000 × g for 10min to remove debris and the supernatant portions collected. The samples were diluted fivefold with 50 mM triethylammonium bicarbonate and BCA assay was carried out to calculate the protein concentration and all samples were normalized by protein amount. Then 250 ug of each sample was digested with Lys-C (Wako) at 1:100 (wt/wt) enzyme-to-protein ratio for 3 h at 37°C. Trypsin was added to a final 1:50 (wt/wt) enzyme-to-protein ratio for overnight digestion at 37°C. To remove the PTS surfactants from the samples, the samples were acidified with trifluoroacetic acid (TFA) to a final concentration of 1% TFA, and ethyl acetate solution was added at 1:1 ratio. The mixture was vortexed for 2 min and then centrifuged at 16,000 × g for 2 min to obtain aqueous and organic phases. The organic phase (top layer) was removed, and the aqueous phase was collected. This step was repeated once more. The samples were dried in a vacuum centrifuge and desalted using Top-Tip C18 tips (Glygen) according to manufacturer’s instructions. The samples were dried completely in a vacuum centrifuge and used for phosphopeptide enrichment using PolyMAC Phosphopeptide Enrichment Kit (Tymora Analytical) according to the manufacturer’s instructions.

The sample was dissolved in 10.5 μL of 0.05% trifluoroacetic acid with 3% (vol/vol) acetonitrile, and 10 μL of each sample was injected into an Ultimate 3000 nano UHPLC system (Thermo Fisher Scientific). Peptides were captured on a 2-cm Acclaim PepMap trap column and separated on a 50-cm column packed with ReproSil Saphir 1.8 μm C18 beads (Dr. Maisch GmbH). The mobile phase buffer consisted of 0.1% formic acid in ultrapure water (buffer A) with an eluting buffer of 0.1% formic acid in 80% (vol/vol) acetonitrile (buffer B) run with a linear 90-min gradient of 6–30% buffer B at flow rate of 300 nL/min. The UHPLC was coupled online with a Q-Exactive HF-X mass spectrometer (Thermo Fisher Scientific). The mass spectrometer was operated in the data-dependent mode, in which a full-scan MS (from m/z 375 to 1,500 with the resolution of 60,000) was followed by MS/MS of the 15 most intense ions (30,000 resolution; normalized collision energy - 28%; automatic gain control target (AGC) - 2E4, maximum injection time - 200 ms; 60 sec exclusion). Minimum precursor mass was set at 350 Da, with the lowest charge state of 2 and the highest of 6. The S/N threshold was set at 1.5 and the minimum peak count of 1. In this dataset, less than 9% of all phosphopeptides had 2 missed cleavages.

### 12. LC-MS Data Processing

The raw files were searched directly against the mouse database with no redundant entries, using Byonic (Protein Metrics) and Sequest search engines loaded into Proteome Discoverer 2.3 software (Thermo Fisher Scientific). The data from the two search engines was combined together. In most cases, the same peptides were identified both. The final data reported is the combination of both search engines, including the identifications reported by only one search engine. MS1 precursor mass tolerance was set at 10 ppm, and MS2 tolerance was set at 20ppm. Search criteria included a static carbamidomethylation of cysteines (+57.0214 Da), and variable modifications of phosphorylation of S, T and Y residues (+79.996 Da), oxidation (+15.9949 Da) on methionine residues and acetylation (+42.011 Da) at N terminus of proteins. Search was performed with full trypsin/P digestion and allowed a maximum of two missed cleavages on the peptides analyzed from the sequence database. The false-discovery rates of proteins and peptides were set at 0.01. All protein and peptide identifications were grouped and any redundant entries were removed. Only unique peptides and unique master proteins were reported.

Total proteomics and phospho-proteomics triplicated data for WT and GS mice treated with haloperidol or saline contained the [log2 intensity value per gene or phospho-peptide normalized to the mean of the run]. Total proteomics data were QCed to i) remove the entries with no gene-name and ii) remove duplicated genes (keeping the replica with the smallest standard error on the fold change haloperidol vs saline). Phospho-proteomics data were only QCed to remove the entries with no gene-name. The number of proteins captured by total proteomics was 3,997 proteins, while the total number of phospho-peptides was 9,399 mapping to a total of 3,010 unique proteins. The fold-change (FC) for the total protein level was calculated for both WT and GS as protein intensity after haloperidol treatment / protein intensity after saline treatment (t-test, two-tailed and homoscedastic). FC = mean(log2(treated)) - mean(log2(saline)). For phospho-proteomics the log2 intensity level of each phospho-peptide was normalized to the log2 intensity level of the total corresponding protein (for each run of saline and haloperidol treatments): Log2 normalized peptide intensity= log2(phospho-peptide) - log2(total protein) before calculating the FC as: peptide intensity after haloperidol treatment / peptide intensity after saline treatment (t-test, two-tailed and homoscedastic). FC = mean(log2(treated)) - mean(log2(saline)). Of note, not for all phospho-peptides a corresponding protein was available in the total proteomics analysis, this reduced the total amount of phospho-peptides that could be analyzed down to 6,787 peptides mapping to a total of 1,641 unique proteins. Threshold for significance was set to p<0.05 (unadjusted p-value) & FC (difference on the log2 intensity values) ≥ |2|. To capture a greater number of phosphorylations for GO-term and kinase-substrate analysis, the statistical cut-off for inclusion was set at uncorrected p > 0.05.

### 13. Pipelines for proteome analysis

GSEA analysis of significantly enriched phosphopeptides for each comparison were analyzed in the listed ontology groups using the enrichR R package to query the Enrichr database^88^

Volcano plots: All quantified phosphopeptides (for proteomic sets) were graphed according to their log2fc and the statistical significance of the change using the ggplot2 R package.

Heatmaps were generated using the ComplexHeatmap R package^89^

### 14. Multicolor fluorescence in situ hybridization and analysis

The smFISH protocol was implemented according to manufacturer guidelines (ACD RNAscope Multiplex Fluorescent Assay v2). Mice were euthanized with carbon dioxide, decapitated, their brains rapidly removed, and placed immediately on foil on dry ice. Brains were then placed in molds containing OCT embedding medium and snap-frozen on dry ice. Embedded brains were sectioned on a cryostat into 20µm sections, mounted onto Superfrost™ slides (and stored at -80°C. Two sections (at ∼-0.1 and –0.34 mm from Bregma) were processed for probe hybridization. All probes and kits were purchased from Advanced Cell Diagnostics, and the chemicals used are described in detail in the resource table. Slides were counterstained with DAPI for 30 sec, cover slipped with mounting medium. Sections were imaged with the Nikon A1 laser microscope system using a 60X 0.75NA objective, to capture 3 z-stack images across 4 channels: 458 (DAPI), 488 (FITC), 561 (TRITC), 638 (Cy5). Projections were obtained using Maximum Intensity Z-projection in ImageJ. Images were analyzed using the CellProfiler4 speckle counting pipeline.

Each field was processed in Cell Profiler^90,60^ to identify nuclei and measure *Drd2/Nr4a1* using the nuclei as seed objects. DAPI channel was enhanced with Enhance Or Suppress Features module, with feature size set at pixel size 30. Identify Primary Objects module was implemented on identified nuclei, object pixel unit was set for Min 5, Max 50, with threshold strategy set as Global and thresholding method set as Otsu. *Drd1* or *Drd2* channel was enhanced with Enhance Or Suppress Features module, with feature size set at pixel size 10. *Nr4a1* or *Arc* channel was enhanced with Enhance Or Suppress Features module, with feature size set at pixel size 5. Identify Primary Objects module was implemented on identified *Drd1* or *Drd2* signal puncta, object pixel unit was set for Min 3, Max30, threshold strategy for Global and thresholding method set as Manual. Nur77 object pixel unit was set for Min 1, Max10, threshold strategy for Global and thresholding method set as Manual. Masked D1 positive nuclei and Masked D2 positive nuclei were generated with RelateObjects. The number of objects were measured with the Measure Object Intensity module.

### 15. Statistics

Group statistical analyses were done using GraphPad Prism 10.1 software (GraphPad, LaJolla, CA). For n sizes, the number of animals and sections are provided. All data are expressed as mean + SEM, or individual plots. Unless otherwise indicated, statistical significance was determined by two-tailed Student’s t-tests for two-group comparisons. For multiple group comparisons, one-way or two-way analysis of variance (ANOVA) tests were used for normally distributed data, followed by post-hoc analyses.

---

**Supplementary Figure 1.**
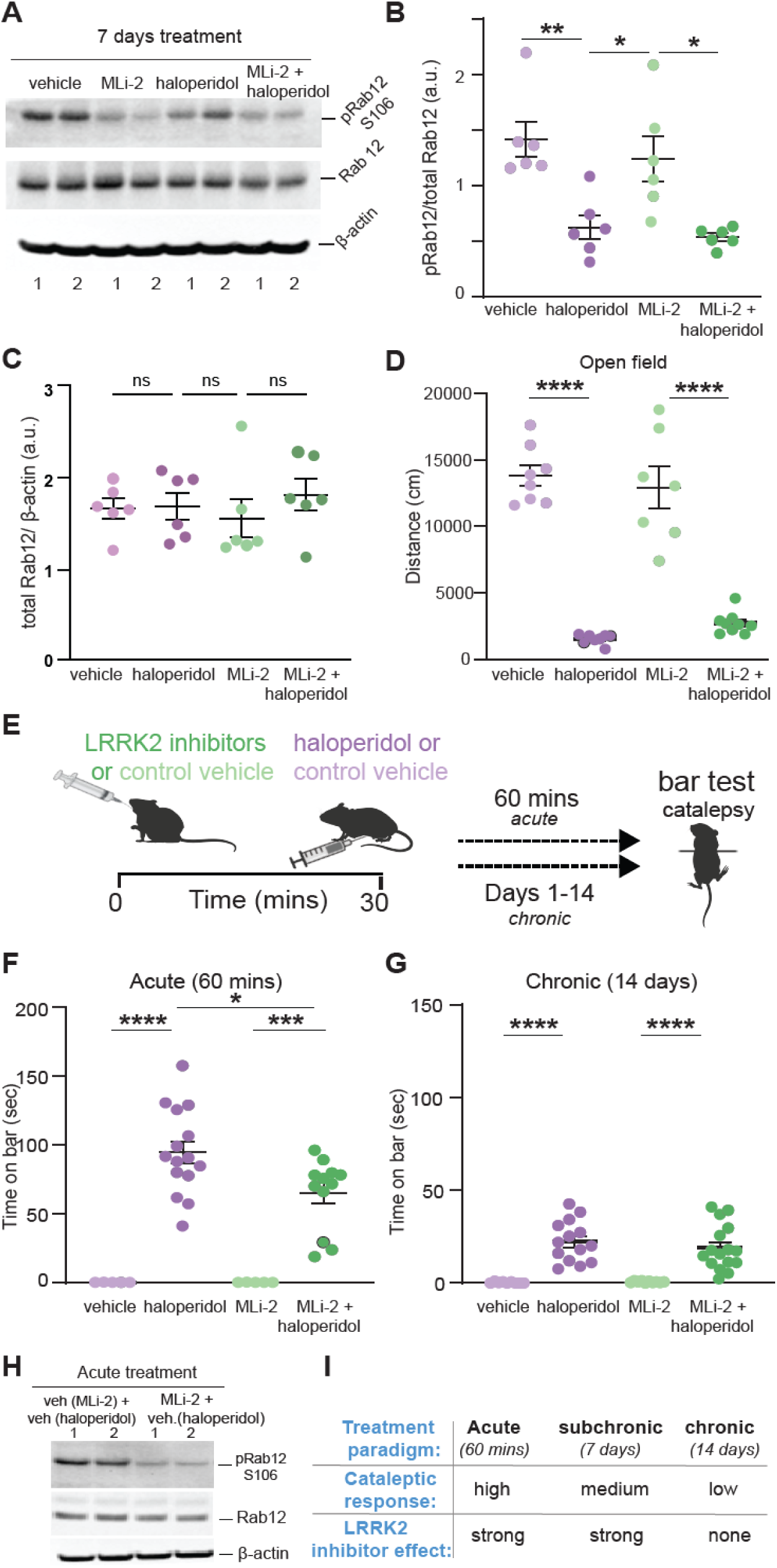
(linked to Figure 1). LRRK2 mediates the effects of haloperidol on movement disruption. **A.** Western blot analysis of striatal extracts from mice across pharmacological treatments for 7 days, probed for pS106 Rab12 (LRRK2 kinase target), total Rab12, and β-actin. **B.** Quantification of p-Rab12 band intensities normalized to total Rab12. n=6 mice **C.** Quantification of total Rab12 band intensities normalized to total β-actin. n=6 mice **D.** Distance traveled in the open field after 7 days of indicated pharmacological manipulations. N= 8, 8, 7, 8 mice, in order of groups presented. **E.** Example of haloperidol and MLi-2 acute and chronic dosing schedule. Catalepsy was assessed 1 hour after the final haloperidol injection. Parts of the schematic were created with BioRender.com. **F.** Catalepsy response of mice treated with haloperidol, MLI-2, or their combination. N=5, 13, 5, 12, in order of groups presented. **G.** Cataleptic response after 14 days administration of haloperidol, MLi-2, or MLi-2 +haloperidol. N=9, 14, 10, 16. **H.** Western blot analysis of striatal extracts from mice treated with MLI-2 or vehicle for 90 mins, as in F, probed for pS106 Rab12 (LRRK2 kinase target), total Rab12, and β-actin. **I.** Table summarizing the magnitude of cataleptic response and the effects of LRRK2 inhibitor in cataleptic response across acute (1 hour), subchronic (7 days), and chronic (14 days) haloperidol treatment paradigms. Data are represented as mean±SEM. Asterisks in B, C, D, F and G denote statistical significance for Tukey’s multiple comparison tests after one-way ANOVA. *p<0.05, **p<0.01, ****p < 0.0001.

**Supplementary Figure 2.**
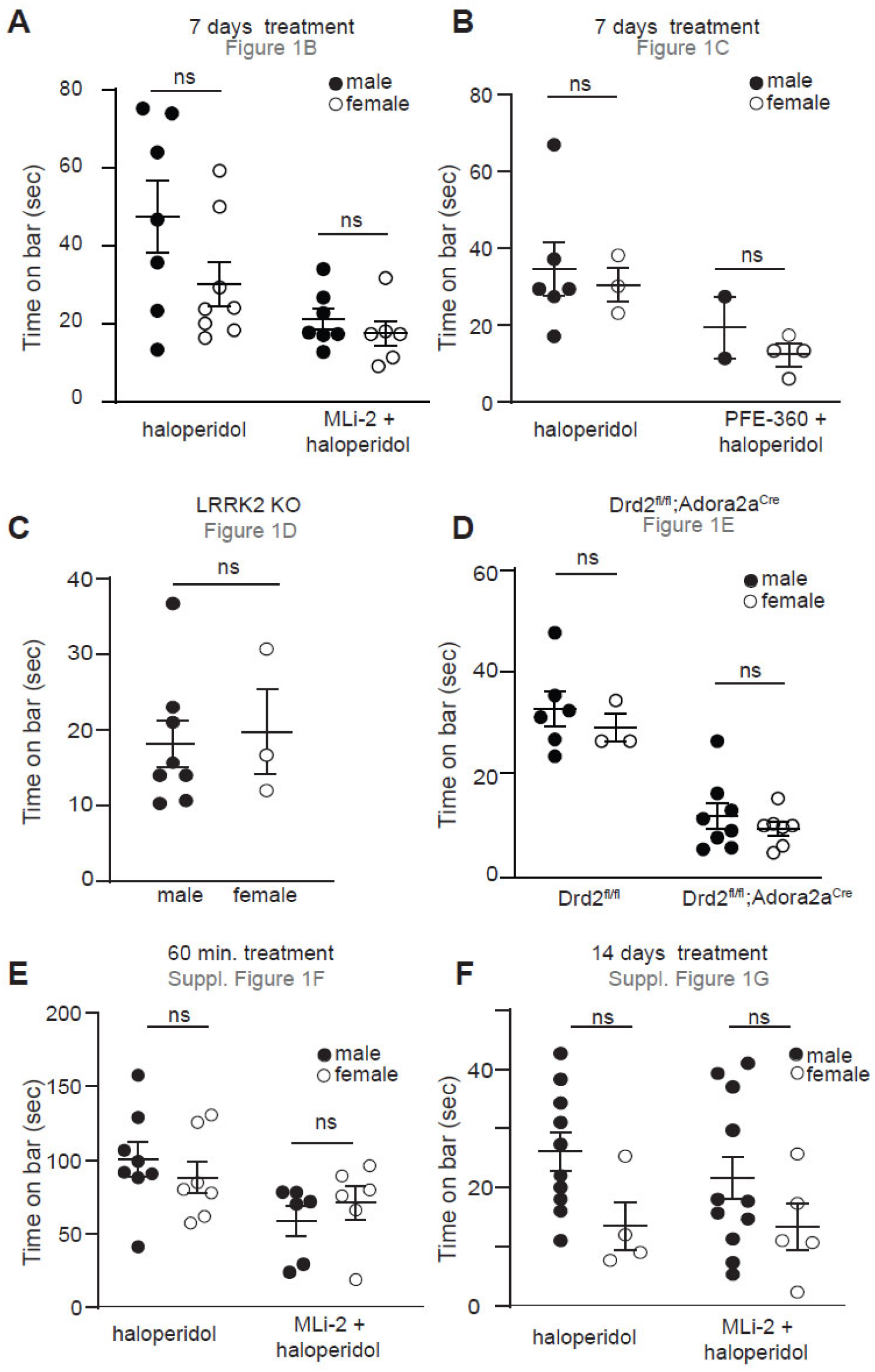
(linked to Figure 1). Haloperidol induced cataleptic behaviors by sex. **A-F**. Summary graph showing the effect of sex across genotypes, pharmacological manipulations, and timepoints for data presented in Figure 1 and Supplementary Figure 1. Relevant figure panels are indicated. Data are represented as mean±SEM. ns, not significant for Tukey’s multiple comparisons test, after two-way ANOVA, except for C (unpaired t-test).

**Supplementary Figure 3.**
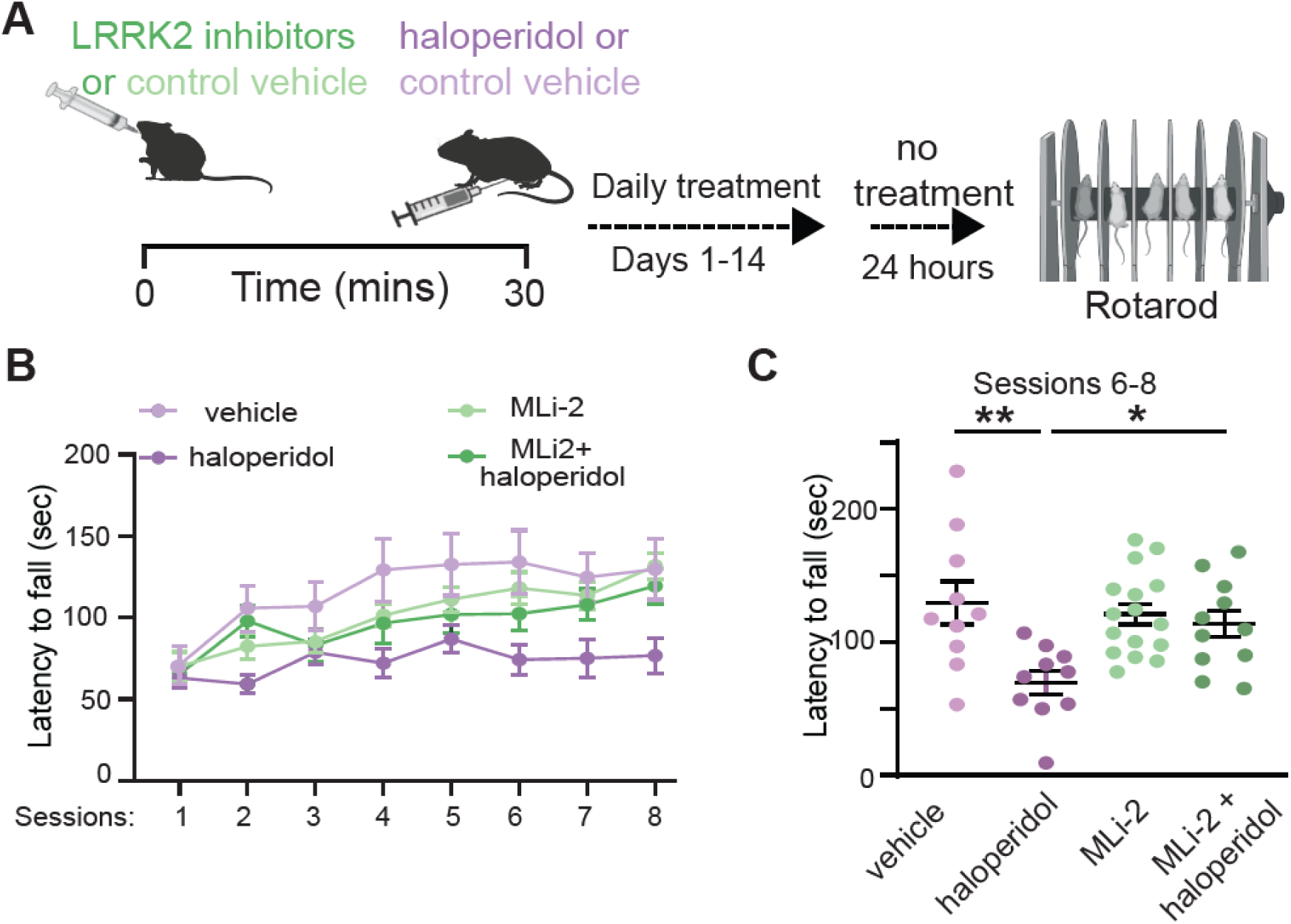
(linked to Figure 1). LRRK2 kinase inhibitors restore haloperidol induced effects on striatal motor learning. **A.** Schematic of the experiment and treatment schedule; it contains schematics created with Biorender.com. **B.** Accelerating rotarod performance (latency to fall) assessed over 8 daily sessions of 5 trials each. Mice received pharmacological compounds, as noted, and were assessed 24 hours after the final haloperidol injection. n=10, 10, 16, 11, in the order are presented. **C.** The average latency to fall in the last 3 sessions of B (Session 6-8). Asterisks show statistical significance for Tukey’s multiple comparison tests after one-way ANOVA; **p< 0.01, *p< 0.05.

**Supplementary Figure 4.**
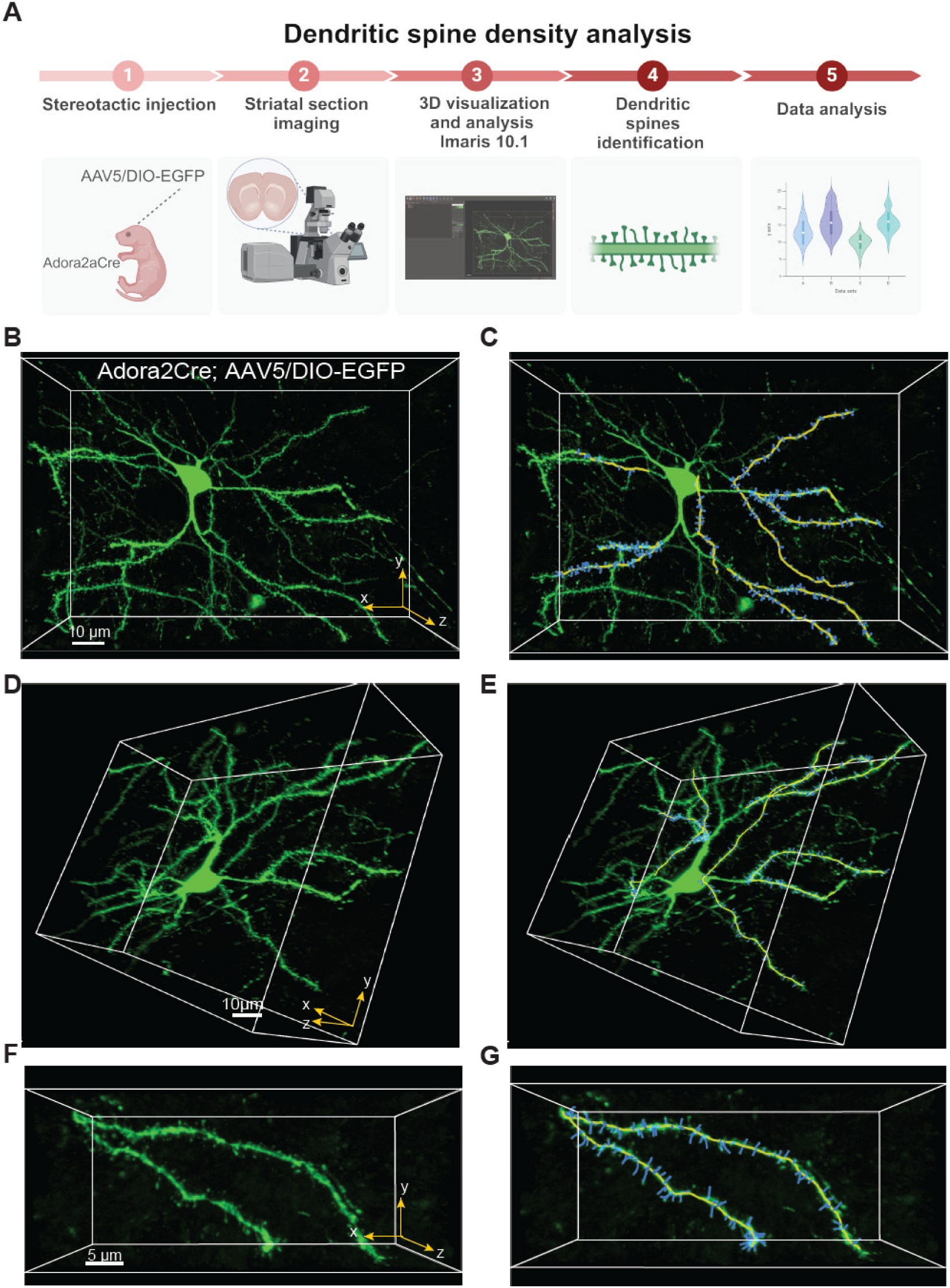
(linked to Figure 2). Dendritic spines density measurement workflow. **A.** Workflow of experimental design for dendritic spine analysis using 3D visualization and analysis with Imaris 10.1. Created with BioRender.com. **B, C.** Representative 3D volume rendering images of an *Adora2a^Cre^* iSPN expressing AAV5/DIO-EGFP, and the corresponding 3D Imaris filament tracer. Scale bar=10 μm **D, E.** A different perspective angle for each x-y-z image in B, C. Scale bar=10 μm **F, G**. Close-ups of dendritic segments from B, C. Scale bar=5 μm

**Supplementary Figure 5.**
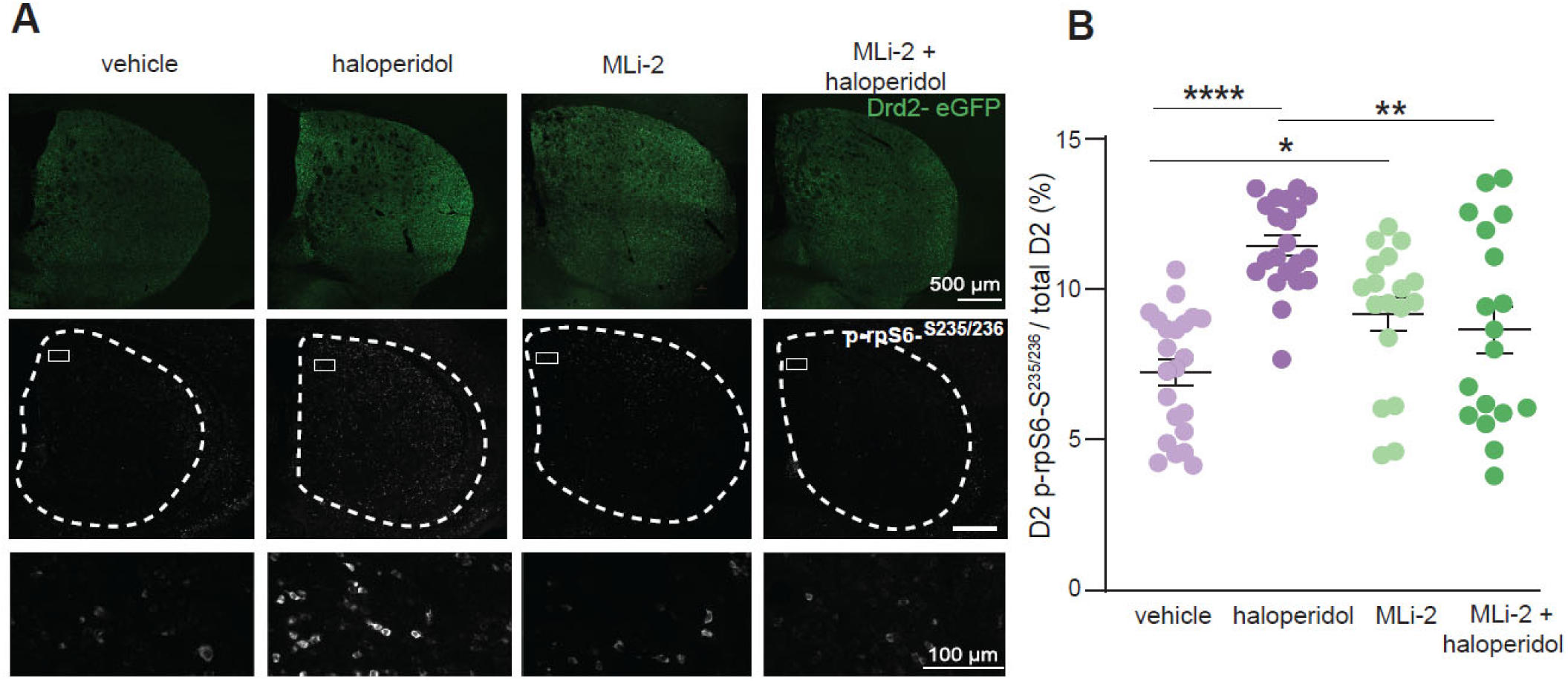
(linked to Figure 3). LRRK2 kinase inhibition restores haloperidol effects on rpS6 signaling in iSPNs. **A.** Representative images showing the distribution of eGFP (top) and p-rpS6 (middle) signal in striatal slices of *Drd2-eGFP* mice treated with vehicle, haloperidol, MLi2, or MLi2+ haloperidol for 7 days. The bottom row shows higher magnification images of p-rpS6 expression. First and second row : scale bar=500 μm; third row: scale bar=100 μm **B.** Ratio of p-rpS6 positive Drd2-eGFP cells to total Drd2-eGFP cells. Data reflect mean±SEM. Asterisks show statistical significance for Tukey’s multiple comparison tests after one-way ANOVA; **p< 0.01, ****p < 0.0001. N=18-22 sections/3-4 mice.

**Supplementary Figure 6.**
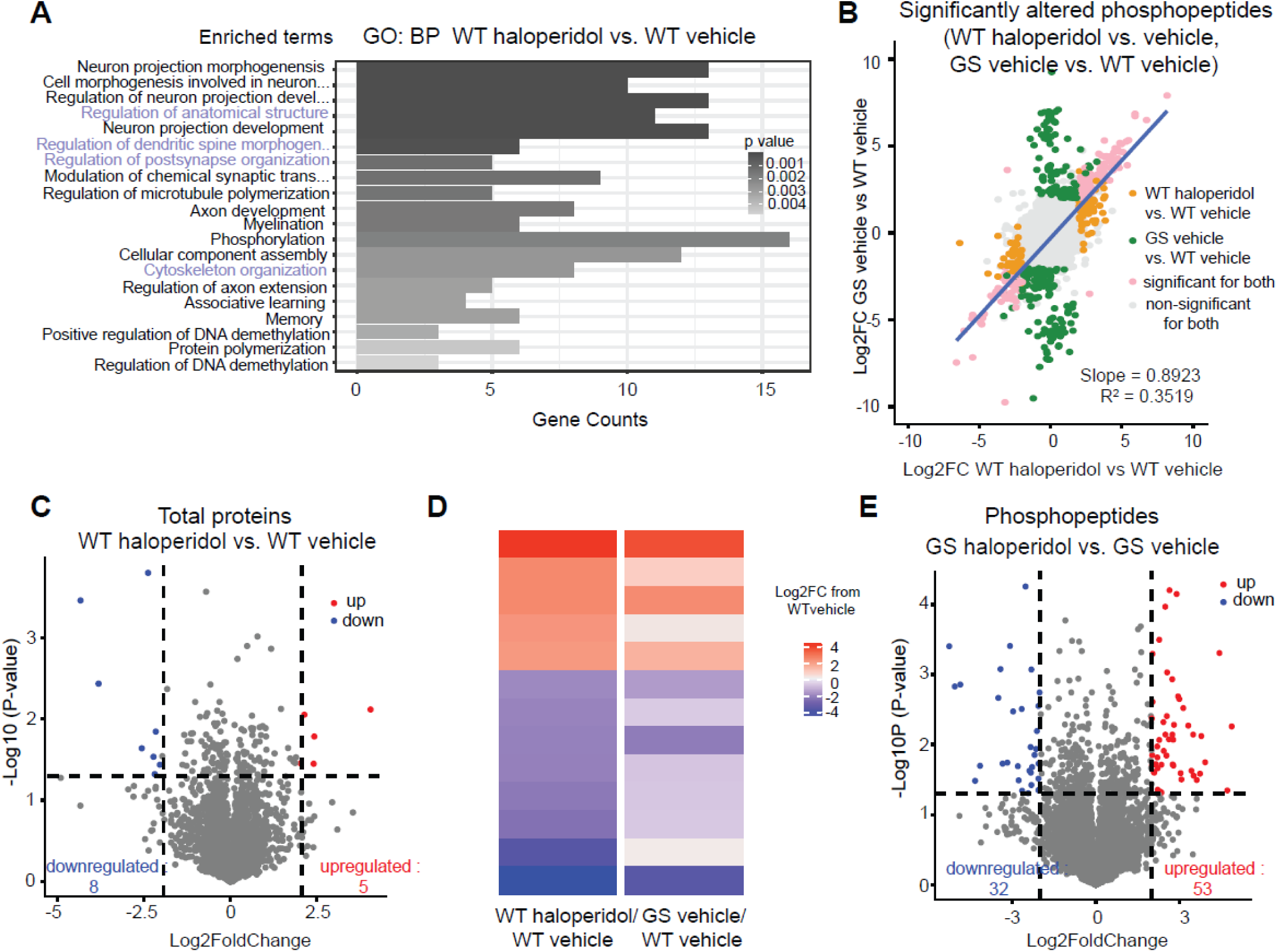
(linked to Figure 3). Proteomic analysis of striatal extracts from haloperidol and vehicle-treated WT and GS mice. **A.** Results of GSEA for the Gene Ontology: Biological Processes gene set on genes with at least one significantly differentially regulated phosphopeptide for the WT haloperidol vs vehicle treated comparison. All pathways displayed are significantly differently regulated (adjusted p-value ≤ 0.05 by Fisher’s test). Length of bars shows the number of genes in the pathway whose phosphostate is differentially regulated, and bars are shaded by adjusted p-value. Highlighted pathways are related to known LRRK2 functions. **B.** Correlation plot for Fig 4E, comparing GS vehicle/WT vehicle to WT haloperidol-vehicle effect size. All points mapped, all significant phosphopeptides (|Log2FC| > 2 and p-value ≤ 0.05 by multiple unpaired t-tests) for either comparison are highlighted and used for correlation. Blue line represents the line of best fit. **C.** Volcano plot of the striatal proteome comparing the haloperidol-vs. vehicle-treated WT mice. Differentially regulated proteins (|Log2FC| > 2 and p-value ≤ 0.05 by multiple unpaired t-tests) are colored in red and blue for up- and downregulated, respectively. **D.** Heatmap of effect size (Log2FC) of either haloperidol treatment or LRRK2-GS for all proteins differentially expressed (|Log2FC| > 2 and p-value ≤ 0.05 multiple unpaired t-tests) compared to the vehicle-treated wild type condition in the wild type haloperidol-vs vehicle-treated comparison, mapped for both the wild type haloperidol-vs vehicle-treated comparison and the GS vs. wild type vehicle-treated comparison. Each bar represents a protein **E.** Volcano plot of relative phosphopeptide for haloperidol- and vehicle-treated LRRK2-GS mutant mice. Phosphopeptides that are differentially regulated (|Log2FC| > 2 and p-value ≤ 0.05 by multiple unpaired t-tests) are colored in red and blue for up and downregulated, respectively.

**Supplementary Figure 7.**
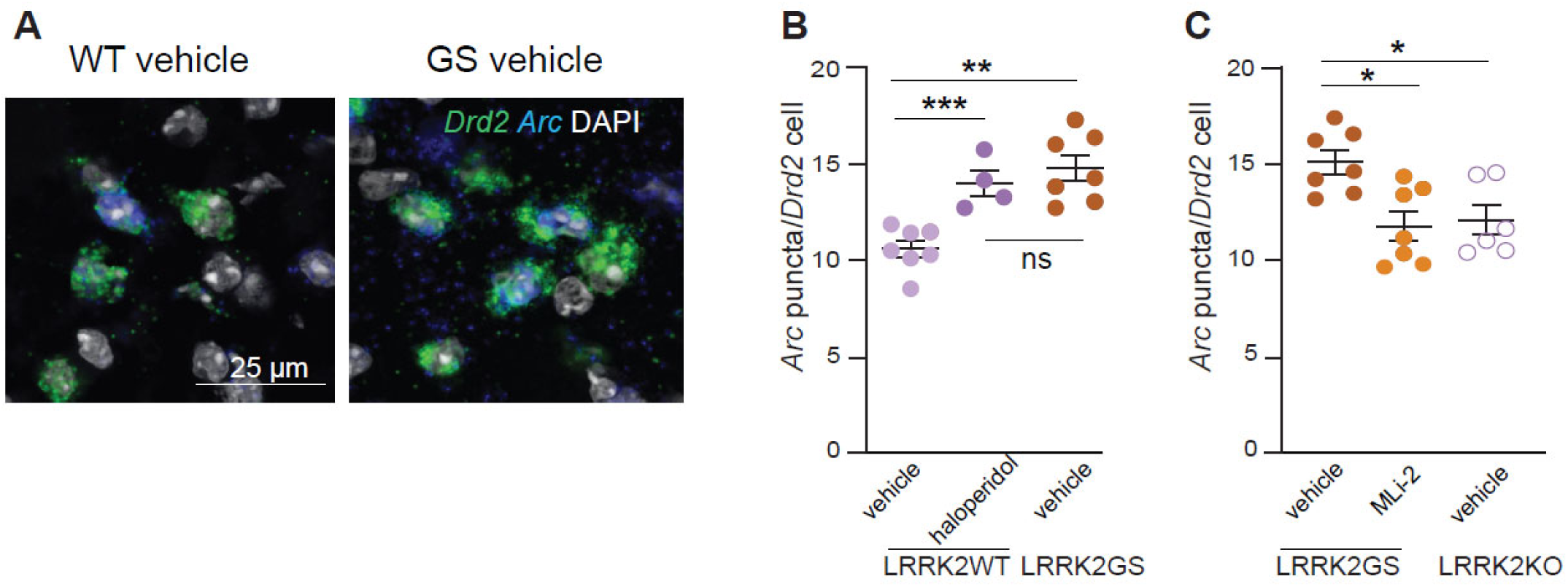
(linked to Figure 4). LRRK2 kinase activity underlies *Arc* increase in indirect pathway SPNs. **A.** Example confocal images of *Arc* gene expression in iSPNs of LRRK2-WT and LRRK2-GS mice. Scale bar=25 μm **B.** Quantification of the number of *Arc puncta* among *Drd2*-positive nuclei. LRRK2-WT mice treated with haloperidol, or vehicle and LRRK2-GS mice treated with vehicle. Each dot represents the average number of *Arc* puncta among *Drd2*-positive nuclei from one striatal section, n=4-7 sections/3-4 mice/group. **C.** Quantification of *Arc* puncta among *Drd2*-positive nuclei in LRRK2-GS mice treated with vehicle, MLi2, and LRRK2-KO mice treated with vehicle for 2 hours. N=6-7 sections/3-4 mice. Data reflect mean±SEM. Asterisks in B, C reflect statistical significance for Tukey post-hoc comparisons after one-way ANOVA. *p < 0.05, **p< 0.01, ***p < 0.001.

